# CHALLENGER: Detecting Copy Number Variants in Challenging Regions Using Whole Genome Sequencing Data

**DOI:** 10.1101/2025.11.23.690083

**Authors:** Mehmet Alper Yilmaz, Ahmet Arda Ceylan, A. Ercument Cicek

**Affiliations:** Department of Computer Engineering, Bilkent University, Ankara, Türkiye

## Abstract

Copy number variation (CNV) detection remains a major challenge in whole-genome sequencing (WGS) data, particularly within repetitive, duplicated, and camouflaged genomic regions where short-read sequencing (srWGS) often fails to produce confident alignments. Although long-read WGS (lrWGS) substantially improves structural variant resolution, its high cost limits widespread adoption, especially in clinical settings. To address these limitations, we introduce CHALLENGER, a masked language modeling–based approach for clinical CNV detection using short-read depth signals over coding regions. While the model uses only short-read data as input, it can make calls typically accessible only with long reads, providing a cost-effective way to obtain information characteristic of both technologies. The model is pre-trained on semi–ground truth calls made on srWGS data and then fine-tuned using (i) lrWGS-derived, (ii) human expert–labeled, and (iii) experimentally validated CNV call sets, enabling it to learn technology- and labeling strategy–specific variant signatures hidden within srWGS profiles and to operate in challenging genomic regions. We show that our short-read–only approach improves the state-of-the-art CNV detection F1-score by 40.8%, while, for the first time, capturing 80.3% of CNVs that can only be detected using long reads in challenging genomic regions. The improvement in F1-score in the set of human experts calls is 70.5% for duplications, and 24.6% for deletions in challenging genes. We also specialize CHALLENGER on paralog genes *SMN1/2, AMY1/2*, and *NPY4R*, and show that it can improve the performance on experimentally validated call sets while being able to make paralog-specific calls in addition to aggregate calls. The CHALLENGER code and model are available at GitHub.

## 1 Introduction

Copy number variations (CNVs), deletions or duplications of large genomic segments, represent a major source of human genetic variation with broad implications for health and disease. Their effects span various biological processes and contribute to neurodegenerative, infectious, autoimmune, metabolic, neuropsychiatric, and developmental disorders [1–4]. In particular, CNVs are estimated to be the basis for up to 15% rare monogenic diseases [5, 6]. Unfortunately, robust detection and functional characterization of CNVs in the clinical setting remain technically challenging and have not yet been fully resolved.

Short-read sequencing platforms remain the foundation of modern genomics because of their high throughput and low per-base cost. Numerous computational tools have been developed to detect of structural variants (SVs) using these technologies [7–17]. However, whole genome sequencing based on short reads (srWGS) continues to face limitations in the assembly or alignment of complex genomic regions, resulting in the so-called dark regions [18]. These regions often arise from high GC content or from duplicated and repetitive DNA elements, such as tandem repeats and gene duplications, that prevent unique read alignment, producing camouflaged regions [19, 20]. Such regions often overlap with genes involved in neurological disorders, including Alzheimer’s disease, autism spectrum disorder, amyotrophic lateral sclerosis (ALS), and spinal muscular atrophy (SMA) [18, 21]. Consequently, short-read sequencing remains limited in its ability to comprehensively detect and interpret CNVs within repetitive or structurally complex genomic regions, restricting the detection of disease-associated variants in climical settings [19, 20].

Advances in whole genome sequencing based on long-read technologies (lrWGS), which sequence thousands to millions of contiguous nucleotides from single DNA molecules, enhance the sensitivity and accuracy of SV discovery in the human genome. Unlike srWGS, lrWGS generates reads that span entire SVs, enabling their direct and unambiguous detection. Long-read sequencing also substantially reduces the number of dark and camouflaged genomic regions that are inaccessible to short reads; for example, Oxford Nanopore Technologies (ONT) data resolve approximately 77% of dark gene body regions [18]. Despite these advantages and advances in lrWGS-based CNV callers [22–28], CNV detection using long reads faces notable limitations, especially in clinical settings. Its sequencing cost remains substantially higher than that of srWGS [29], and its relatively low coverage reduces sensitivity for large CNVs, which are often better detected by read depth-based analyzes [30]. Indeed, some lrWGS assembly methods capture only a subset of large (>5 kb) CNVs detected by srWGS, highlighting persistent blind spots in long-read assemblies [30].

Integration of multiple sequencing technologies remains essential to achieve an accurate and more complete characterization of CNVs in the human genome. However, simultaneously using multiple technologies, such as using both short- and long-sequencing data at the same time, is not feasible due to the multiplying costs and are only available in certain research resources such as 1000 Genomes Project. Such datasets provide a valuable resource for machine learning models which can learn a mapping between the output of the complementing technologies. This is a promising direction for researchers to use a single, and preferably cheaper technology and to exploit the benefits of others without performing sequencing but through modeling. Previous work have used this idea to increase the CNV calling accuracy (i) on whole exome sequencing (WES) data [10, 13], and (ii) on ancient WGS data [31]. Unfortunately, none of the available WGS-based CNV detection methods (either srWGS-based or lrWGS-based) can take advantage of both technologies at the same time and they cannot model one using the other. Moreover, no existing WGS-based CNV caller can mimic calls in highly curated, multi-technology CNV variant call sets such as those from Chaisson et al. [32] which are considered ground truth. These callsets capture an integration of long-read, short-read, optical mapping, and Hi-C data, representing a gold standard that current single-technology callers cannot model. The inability to infer such multi-modal variant patterns from a single data type remains one of the biggest obstacles in clinical CNV calling.

Here, we present CHALLENGER, a transformer-based deep learning framework which for gene-level CNV detection using srWGS data. For the first time, our model which uses srWGS as input, can pinpoint CNVs that can only be detected using orthogonal technologies such as lrWGS or by human experts. It can also be optimized for accurate CNV calling in the challenging regions of the genome. The model processes read-depth signals for each gene through a cascade of convolutional and transformer encoder blocks, and incorporates a trainable token to capture genomic context. We pre-train the model using CNV calls derived from hundreds of WGS samples from the 1000 Genomes Project [33] annotated with the DRAGEN [12]. This pre-training enables the model to learn robust relationships between read-depth patterns and semi-ground truth CNV events in srWGS data. To extend its capability to multi-technology CNV detection, we fine-tune the model using (i) long read-based CNV calls alongside short read-based calls, (ii) high-confidence CNVs validated by human experts and multiple orthogonal technologies, and (iii) experimentally validated gene-specific CNV call sets. These steps allow CHALLENGER to capture cross-technology patterns in short read-based read depth signals associated with CNVs. Extensive benchmarking demonstrates that CHALLENGER achieves state-of-the-art performance in CNV detection across multiple sequencing modalities. We show that our approach improves the state-of-the-art CNV detection F1-score in challenging genes to 70.7% (an improvement of 40.8% over the next best method), while, for the first time, capturing 80.3% of CNVs that can only be detected using long reads. Moreover, our F1-score across all genes is 64.5% (an improvement of 28.1% over the next best method). We also observe substantial improvement in F1-score in detecting human experts calls in the set of challenging genes where we obtain an F1-score of 87.4% for duplications (an improvement of 70.5% over the next best method) and 79.7% for deletions (an improvement of 24.6% over the next best method). We also specialize CHALLENGER to make accurate calls on paralog genes which are of critical clinical importance (*SMN1/2, AMY1/2, and NPY4R*), and show that the model improves the overall performance on these genes while enabling making paralog-specific and aggregate calls. The CHALLENGER code and model are available at https://github.com/ciceklab/CHALLENGER.

## 2 Results

### 2.1 Model Overview

The model has a transformer-based architecture that adapts RoBERTa [34] for copy number variation detection using srWGS. Figure 1 shows the system model. CHALLENGER takes the read-depth signal on a given gene region as input, which is averaged using 50 bps bins to reduce noise and computational complexity. To ensure consistent input dimensionality, each gene region is represented as a fixed length vector (1000). For longer regions, the bin size is adaptively increased, whereas shorter regions are padded. After normalization, the read-depth sequence is subsequently processed by a two-layer convolutional neural network (CNN), which projects the signal into a higher-dimensional latent space to generate read depth embeddings. In addition to read depth embeddings, CHALLENGER generates three types of embeddings: (i) *Gene embeddings* that encode gene-specific characteristics, (ii) *CNV embeddings* that represent the CNV labels corresponding to read depth embeddings, and (iii) *Positional embeddings* that capture the relative position of each token within the sequence. During pre-training, a portion of the CNV embeddings is masked, with the masking ratio gradually increased in later training stages; in fine-tuning, all CNV embeddings are fully masked to promote faster learning and stronger contextual understanding. Finally, all embeddings (read-depth, gene, CNV, and positional) are summed element-wise, and the resulting representation is passed through three cascaded transformer layers. The first element of the final hidden state of the transformer layer, corresponding to the gene-level classification token, is used to predict the CNV event for the entire gene through fully connected layers. The remaining token-level outputs are passed to a token-level classification layer (used only during pre-training) to predict CNV labels for each masked CNV token. Details about each step are provided in the Methods Section. To justify the need for such a complex architecture, we conducted an ablation study to examine the effects of removing (i) transformer layers, (ii) gene embeddings, and (iii) positional embeddings. We find that removing each component substantially reduces model performance, showing their importance in capturing gene-specific contextual patterns and long-range dependencies within read-depth signals. (See Supplementary Table 23).

**Fig. 1.**
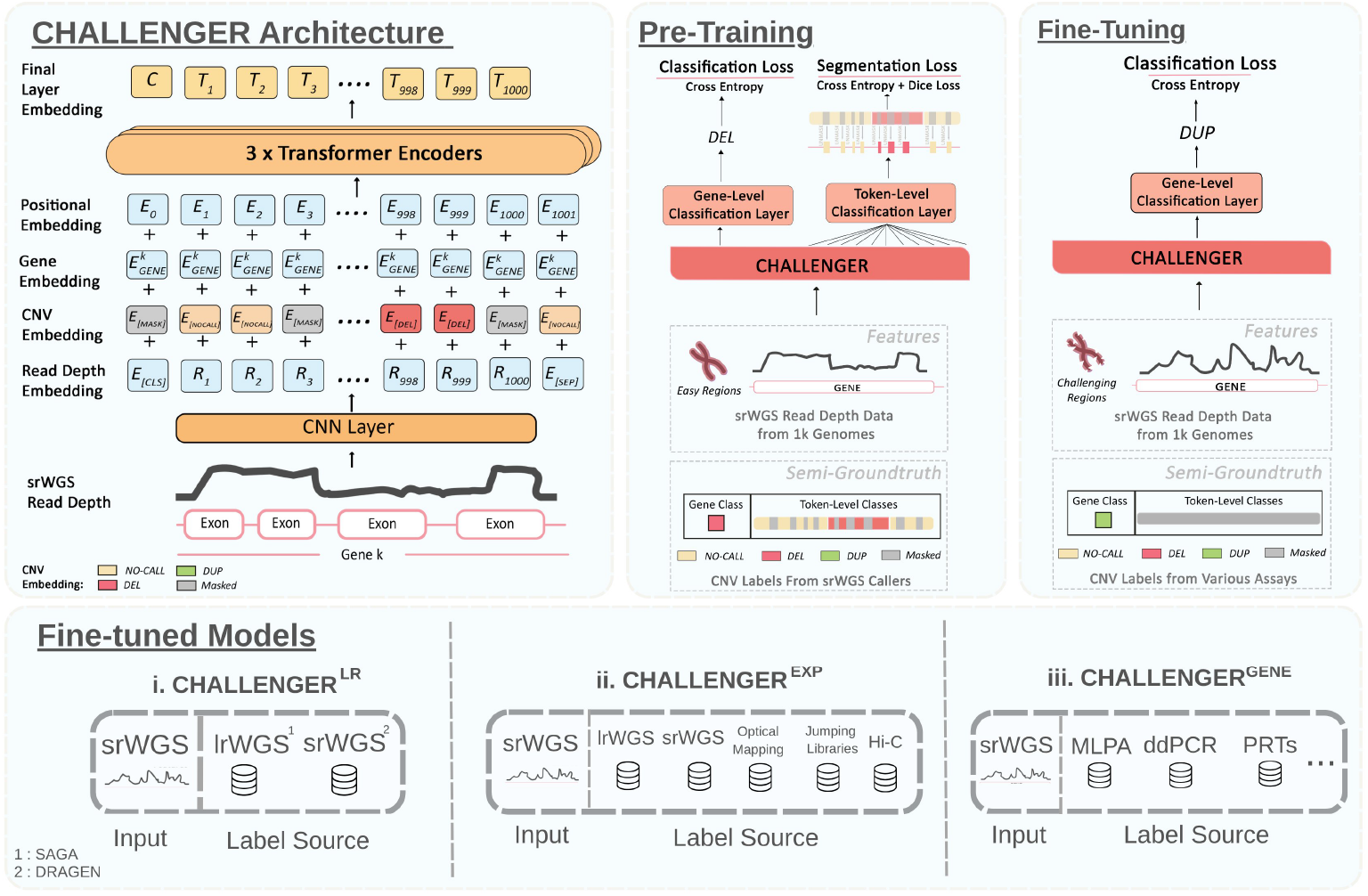
The system model of CHALLENGER. **Architecture**. The model embeds the normalized read-depth signal of each gene using two sequential 1D convolutional layers, each followed by batch normalization and ReLU activation, producing a high-dimensional read-depth embedding. A trainable classification token and separator tokens are appended to the sequence, which is then summed with (i) CNV embeddings that indicate the underlying copy-number state (NOCALL, DEL, DUP, or MASK), (ii) gene embeddings that encode gene-specific read-depth characteristics, and (iii) positional embeddings. The combined representation is fed into three transformer encoder blocks to produce contextualized token and gene-level features. **Pre-training**. During pre-training, CHALLENGER jointly learns gene-level and token-level CNV prediction using DRAGEN-derived labels, with progressively increased masking of CNV embeddings. **Fine-tuning**. In fine-tuning, only the gene-level head is used, allowing adaptation to diverse CNV resources. **Fine-tuned Models**. We present 3 specialized fine-tuned variants: CHALLENGER^LR^ (fine-tuned on short-read and long-read–derived CNVs), CHALLENGER^EXP^ (trained on multi-technology-based and human expert-curated CNV callsets), and CHALLENGER^GENE^ (gene-specific models trained on experimentally validated CNVs available for *SMN1/2, AMY1/2*, and *NPY4R*).

### 2.2 Evaluation criteria

For each gene, CHALLENGER makes a deletion, duplication, or no-call decision. We evaluate its performance at the gene level using precision, recall and the F1 score. We assign semi-ground truth or ground truth CNV labels to genes by intersecting the detected CNV regions in the call set with the corresponding gene regions. Calls in paralogous genes are evaluated both individually and also in an aggregated manner (i.e., making a single call for both copies), as also done in the literature [35]. We consider CNVs longer than 1 kb in our evaluation. To assess performance in challenging regions, we use the dark and camouflaged gene list based on Illumina 100-bp sequencing data reported by Ebbert et al. [18]. Regions are defined as dark-by-depth when ≤5 reads are aligned and dark-by-mapping-quality when ≥90% of reads have MAPQ *<* 10 [18]. The list includes 3,804 genes in total.

### 2.3 CHALLENGER^LR^ captures both srWGS specific and lrWGS specific CNVs on challenging genes and genome-wide

To fine-tune the model for identifying CNVs derived from long-read whole-genome sequencing (lrWGS) data, we use the CNV callset generated by Schloissnig et al. [36]. The dataset includes 1,019 lrWGS samples sequenced using ONT, together with CNV calls generated using the SAGA framework, which integrates consensus calls from DELLY [28] and Sniffles [22] and provides a golden ground truth CNV callset (see Section 4.1 for details). We use 230 randomly selected samples for fine-tuning and 30 samples for testing. For both training and test sets, we combine the lrWGS-based CNV calls with the DRAGEN-based CNV calls obtained on matched srWGS samples. The details of the sources of CNV labels for the fine-tuning and test sets are shown in Supplementary Table 25 and 26, respectively. This fine-tuned model is denoted as CHALLENGER^LR^.

Since dark and camouflaged genomic regions present a challenge for reliable detection of CNVs on srWGS data, we evaluated CHALLENGER^LR^ and other benchmarked tools specifically for genes located within these regions. The distribution of CNVs captured by each technology across these regions in the test set is shown in Figure 2A. Note that the number of detected deletions is roughly seven times greater than the number of duplications. CNV calls from srWGS and lrWGS overlap by only 0.2% for duplications and 6.1% for deletions, underscoring the importance of integrating both technologies. We compare the performance of CHALLENGER^LR^ with 5 srWGS-based and 3 lrWGS-based callers.

**Fig. 2.**
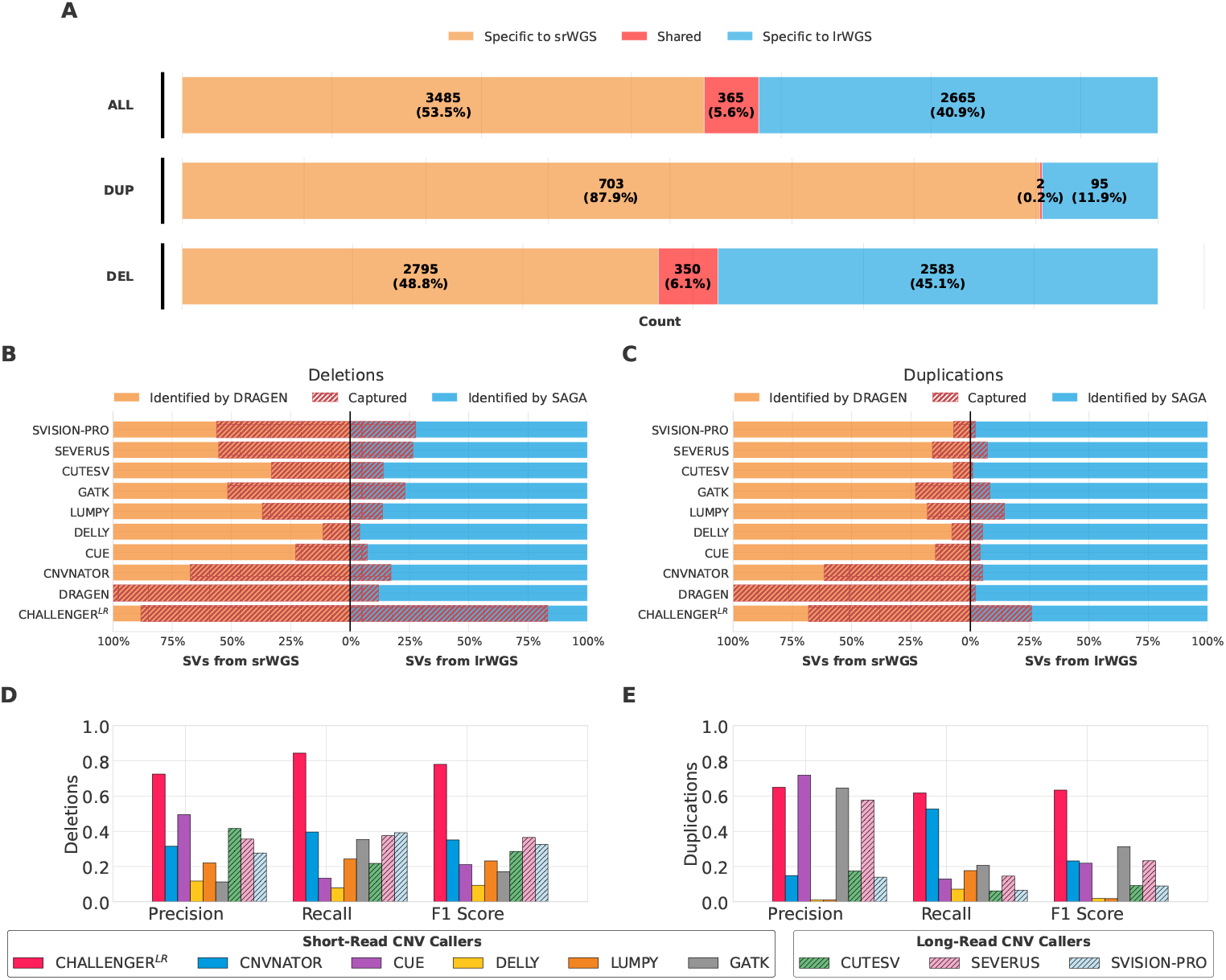
Evaluation of CHALLENGER^LR^ in capturing long-read, and short-read specific CNVs in challenging genomic regions (see Supplementary Figure 1 for genome-wide performance comparison). **(A)** Overlap between CNV calls derived from short-read WGS (srWGS; DRAGEN) and long-read WGS (lrWGS; SAGA) datasets. **(B-C)** Comparative recall of the model and existing tools for CNVs detected in srWGS and lrWGS datasets, shown separately for deletions (B) and duplications (C). **(D-E)** Comparison of precision, recall, and F1-score across methods for deletions (D) and duplications (E). For srWGS-based calls, DRAGEN-based calls are treated as semi-ground truth. For this reason, DRAGEN has 100% recall on srWGS-based CNV calls.

For each method, we show the percentage of CNVs captured from each technology is shown in Figure 2B for deletions and in Figure 2C for duplications with corresponding numeric values available in Supplementary Table 1. Our model achieves the highest recall for both long-read and short-read CNVs. For long-read–based deletions and duplications, it improves recall by 55.7% over SVision-Pro and 11.4% over Lumpy, the next-best methods. For short-read–based CNVs, it outperforms the second-best method, CNVnator, by 20.8% and 6.6%, for deletions and duplications, respectively. For the first time, our model captures 80.3% of CNVs that can only be detected using long reads. Note that in the analysis of srWGS-based calls, DRAGEN-based calls are treated as semi-ground truth. For this reason, DRAGEN has 100% recall on srWGS-based CNV calls. It should be noted that, despite using srWGS data, CHALLENGER^LR^ is capable of capturing more calls based on lrWGS than all callers based on lrWGS. This is because the model has been taught to associate the read depth signal with the consensus call set identified in the SAGA framework which can be considered a golden call set by incorporating multiple tools and manual refinement. The general performance comparisons are shown in Figures 2D and 2E, which show that our model not only has a high recall, but is also precise. In all evaluations, the model consistently shows superior performance, It improves the state-of-the-art CNV detection F1-score in challenging genes to 70.7%. This corresponds to an F1-score improvement of 41.5% over Severus for deletions and 32.1% over GATK for duplications, the second-best methods in each category. We also present the precision and recall results in Supplementary Tables 2 along with the corresponding confusion matrices in Supplementary Table 3.

We extend the same analysis from the list of challenging genes to all genes and observe the exact same trend in challenging regions. The distribution of CNVs captured by each technology in the test set is shown in Supplementary Figure 1A. For each method, the percentage of CNVs captured from each technology is shown in Supplementary Figures 1B and 1C, for deletions and duplications, respectively with the corresponding numeric values provided in Supplementary Table 4. We observe a similar trend genome-wide. The model achieves the highest recall for long-read CNVs, with recalls of 79.2% for deletions and 56.6% for duplications, and also the highest recall for short-read deletions at 89.9%, while its recall for short-read duplications is the second highest at 42.9%. The precision and F1-score results are shown in Supplementary Figures 1D and 1E, as well as in Supplementary Table 5, with the corresponding confusion matrices provided in Supplementary Table 6. We observe F1-score improvements of 37.9% and 12.4% over the second-best methods for deletions (CNVnator) and duplications (GATK), respectively.

### 2.4 Learning from Multi-Technology and Human Expert CNV Calls Expands the Spectrum of Detectable Variants

To improve our model’s ability to detect a wide range of human genetic variation, we fine-tuned it using the structural variant dataset published by Chaisson et al. [32] which is considered as a human expert-labeled golden ground truth callset. This set includes CNVs from nine individuals analyzed by human experts using several complementary technologies, including long-read sequencing, short-read sequencing, Hi-C, strand-specific sequencing, optical mapping, and multiple variant calling methods. Of the eight available samples, six were used for fine-tuning and three were reserved for testing. The resulting fine-tuned model is denoted as CHALLENGER^EXP^.

Figures 3A and 3B show the gene-level performance of the evaluated tools in challenging genomic regions. Detailed performance scores are provided in Supplementary Table 7, along with corresponding confusion matrix in Supplementary Table 8. CHALLENGER^EXP^ consistently outperformed all other methods in both deletion and duplication detection, achieving F1-scores of 79.7% and 87.4%, corresponding to improvements of 24.6% over Severus and 70.5% over CNVnator, the next-best performing tools. These results show that CHALLENGER^EXP^ effectively learns complex, technology-specific signal patterns present in short-read WGS data. By incorporating features that capture the characteristic biases and signal properties of each sequencing technology, CHALLENGER^EXP^ can detect CNVs often missed by conventional algorithms, thereby expanding the range of variants that can be reliably identified. We observe the same trend in all genes. Figures 3C and 3D show the precision, recall, and F1-score performances of all methods throughout the genome. The corresponding performance metrics are provided in Supplementary Table 9, and the confusion matrices are shown in Supplementary Table 10. CHALLENGER^EXP^ results in F1-score improvements of 22.4% and 74.3% over second-best methods for deletion (Severus) and duplication (CNVnator) CNV detection, respectively.

**Fig. 3.**
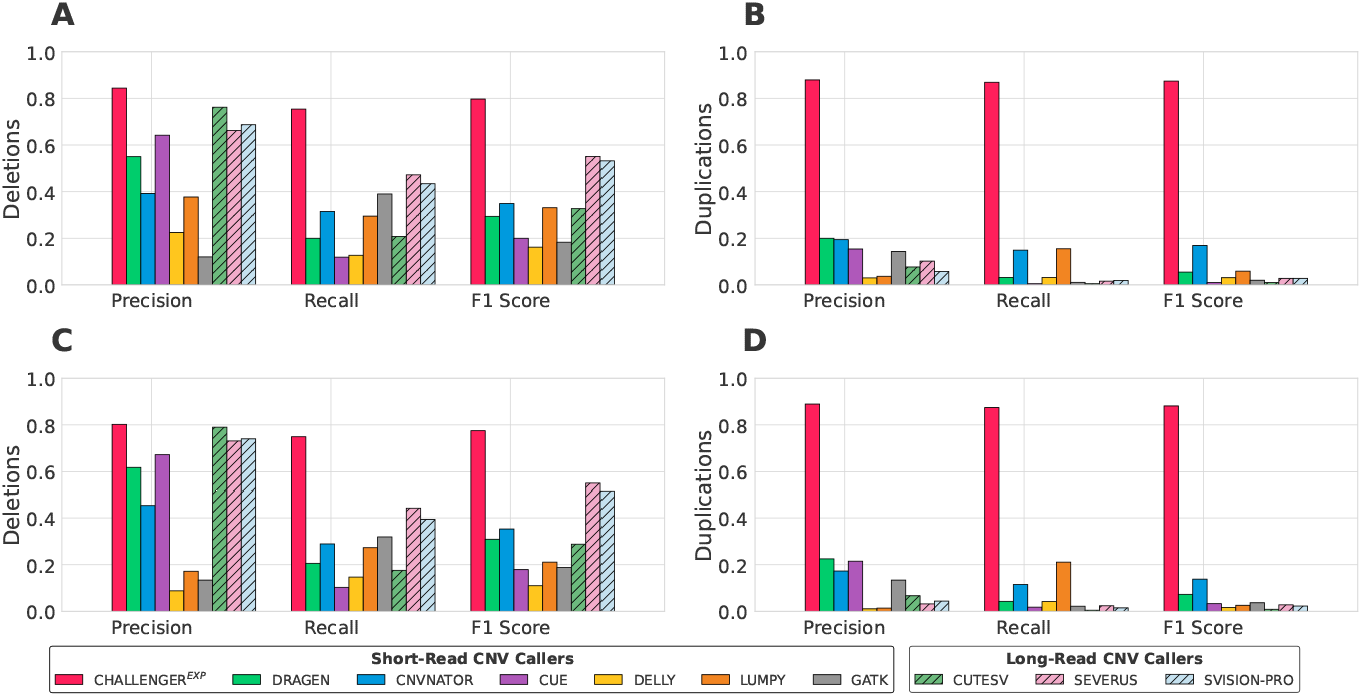
Benchmark of precision, recall, and F1 scores for CNV detection algorithms evaluated on the Chaisson et al. golden CNV call set (**A** and **B**) in challenging genomic regions and (**C** and **D**) for all genes. (**A** and **C**): Deletion performance, (**B** and **D**): Duplication performance.

### 2.5 CHALLENGER can be fine-tuned on experimentally validated CNV call sets to function as a dedicated gene-specific caller

CHALLENGER can also be customized as a gene-specific caller to detect experimentally validated CNVs, which are particularly important in duplicated genomic regions where standard methods often struggle. To achieve this, we fine-tune our base model using experimentally validated CNV call sets for 4 duplicated genes. Specifically, we incorporate CNV events detected with Multiplex Ligation-dependent Probe Amplification (MLPA) for *SMN1/2* [37], droplet digital PCR (ddPCR) for *NPY4RA/B* [38], and paralogue ratio tests (PRTs), duplication junction PCR, and microsatellite assays for *AMY1A/B/C* and *AMY2A* [39]. In our analysis, *AMY1, NPY4R*, and *SMN* are used for aggregate CNV detection, while *SMN1, SMN2*, and *AMY2A* are used for paralog-specific CNV detection. *AMY1* and *NPY4R* are excluded from paralog CNV detection because the corresponding studies report only aggregate copy numbers for these genes. In total, we use 103 samples for fine-tuning and 96 samples for testing (listed in Supplementary Tables 28-30). We denote the variant of CHALLENGER, which is tailored per each gene as described above, as CHALLENGER^GENE^.

We compare our fine-tuned model CHALLENGER^GENE^ with CNV callers that are developed to specifically target paralog genes, Parascopy [35], QuicK-mer2 [40], and SMNCopyNumberCaller (SMNCNC) [37] in both aggregate CNV detection and paralog CNV detection (See Table 1). Supplementary Tables 11–22 report the detailed performance scores and associated confusion matrices for all genes. CHALLENGER^GENE^ achieves the highest F1 scores in 9 out of 11 comparisons. Only in detecting paralog-specific deletion calls for *SMN1*, SMNCopyNumberCaller achieves the best score. Although other tools match CHALLENGER^GENE^’s performance in certain analyzes, they underperform in others. For example, Parascopy cannot make paralog-specific calls by design. QuicK-mer2 performs well in *SMN1/2* aggregate CNV detection, but cannot distinguish paralog-specific calls in *SMN1* and *SMN2* as well. As the name implies, SMNCopyNumberCaller can only call for the *SMN1* and *SMN2* genes. These results show that CHALLENGER^GENE^ can be used effectively for the reliable detection of CNVs in duplicate regions.

**Table 1.**
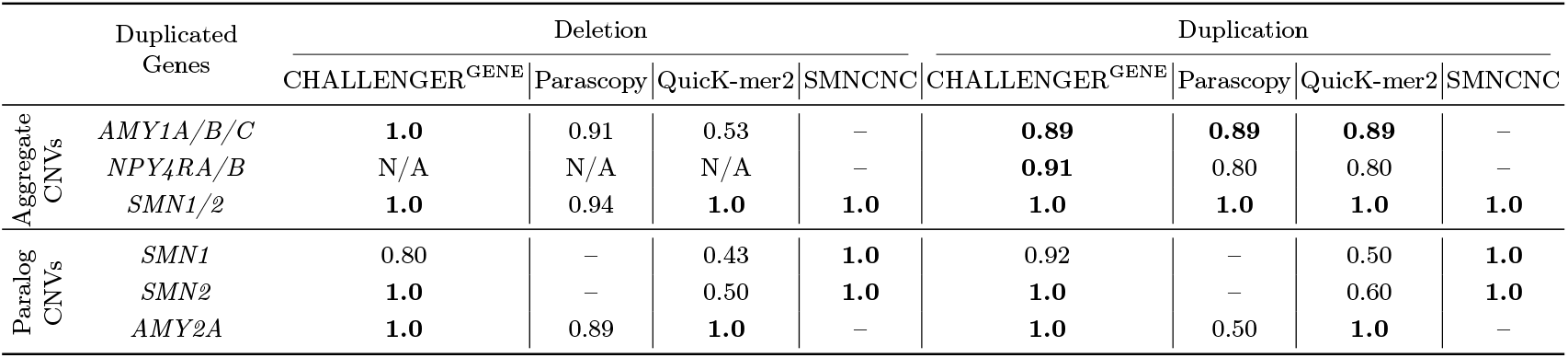
Comparison of F1 scores for detection of experimentally validated CNVs in duplicated genes. Vertical labels denote CNV detection task groups (Aggregate CNV detection and Paralog CNV detection). In the former a single decision is made for the genes in the group and in the latter, gene-specific calls are made. SMNCNC is the abbreviation for SMNCopyNumberCaller. N/A indicates there are no ground truth calls for that gene of that type. – means that Parascopy or SMNCopyNumberCaller cannot make CNV calls by design. The former cannot make paralog-specific calls, the latter cannot make calls for any gene other than *SMN1/2*.

In Figure 4, we compare the IGV visualizations of two samples NA18745 and NA10847 over the *NPY4RA* and *NPY4RB* paralog genes. The left panel shows the NA18745 who has a duplication event, while the right panel shows NA10847 who does not have any CNVs (e.g.,no-call). Both events are validated by ddPCR at the aggregate level. Despite this confirmation, the duplication signal is subtle and difficult to observe or distinguish from NA10847 visually. The read-depth fluctuations are shallow, noisy, and do not form a canonical duplication signature. Consistent with this ambiguity, both Parascopy and QuicK-mer2 cannot identify the duplication for this sample. In contrast, CHALLENGER^GENE^ accurately distinguishes the duplication, and no-call events, even in the presence of weak coverage cues which is powered by teaching the model the relationship between the read depth signature in srWGS data and ddPCR-based CNV labels.

**Fig. 4.**
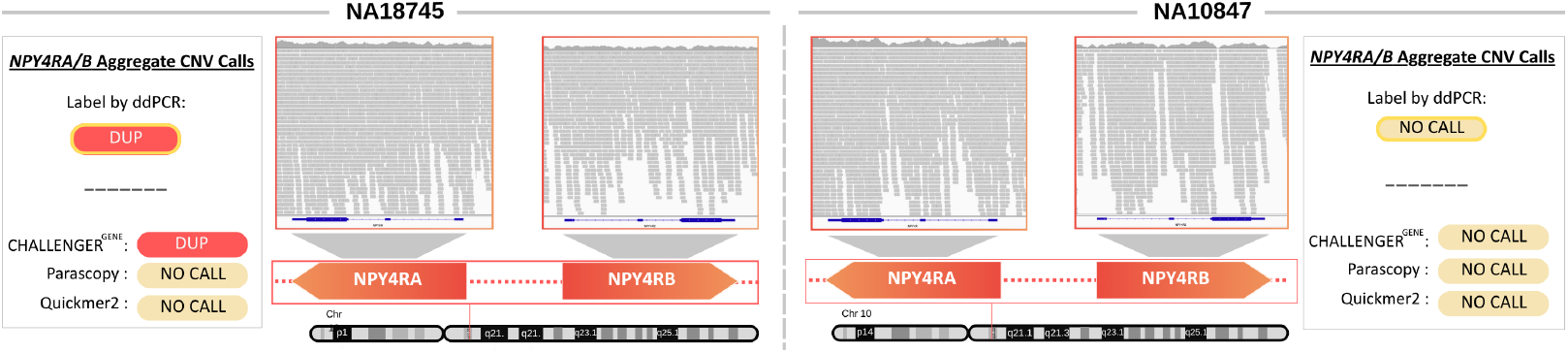
IGV visualizations of the *NPY4RA* and *NPY4RB* paralog gene regions for NA18745 and NA10847 samples. The left panel shows a ddPCR-validated duplication event, and the right panel shows again a ddPCR-validated no-call. Neither Parascopy nor QuicK-mer2 can detect the CNV event in this region, whereas CHALLENGER^GENE^ accurately distinguishes the duplication and no-call events by associating the read depth signal with the experimental labeling technologies via fine-tuning.

## 3 Discussion

In this study, we present CHALLENGER, a deep learning framework designed to detect copy number variants (CNVs) in challenging genes using srWGS data. Using a masked language modeling strategy and bidirectional transformer-based architecture, CHALLENGER effectively learns cross-technology, human expert, and experimentally validated CNV call patterns in srWGS read-depth signal that are typically not identifiable using only short reads. Through extensive fine-tuning on datasets derived from long-read, expert-labeled, and experimentally validated call sets, the model achieves superior performance genome-wide compared to state-of-the-art CNV detection tools which can be srWGS-based, lrWGS-based, or tailored towards duplicated genes. We observe the advantage especially in repetitive and camouflaged genomic regions. CHALLENGER thus represents a significant step towards achieving high-resolution CNV detection using cost-effective short-read sequencing technologies.

An important concern is whether the model’s superior performance arise from label fixation, that is memorizing the label distributions present in the fine-tuning sets rather than learning meaningful cross-technology patterns. Although such overfitting is a legitimate risk for models trained with limited or modality-specific labels, our results indicate that CHALLENGER does not rely on label memorization. Instead, the model can make zero-shot calls. Notably, CHALLENGER^LR^ achieves 100% F1 score in 50 genes for which it encounters no CNV events at all during training or fine-tuning, demonstrating its ability to generalize beyond the observed label space. Moreover, in 189 genes where the allele frequency of CNVs in the training and fine-tuning sets is below 1%, the model again reaches 100% F1 score on the test set (see Supplementary Tables 31-34 for gene lists). These findings indicate that CHALLENGER^LR^ internalizes latent CNV signatures associated with each labeling technology during fine-tuning and can successfully apply these learned representations to short-read input profiles, rather than relying on fixed or memorized label patterns.

Despite these promising results, some limitations remain. A major limitation arises from the unique characteristics of challenging genes, such as high sequence similarity among paralogs, extreme GC content, and complex duplication structures. These properties necessitate the use of gene-specific embeddings to capture gene-level contextual information, ensuring that the model correctly interprets subtle read-depth fluctuations. While this strategy markedly enhances accuracy in those regions, it also constrains the model’s ability to generalize to other regions and comes with the cost of fine-tuning a model per gene. While the number of such genes is not big and this is doable, the model is constrained with the availability of high-quality experimentally-validated CNV labels for duplicated genes. For this reason, us and others have showed results on a small number of examples. We expect the model’s performance and generalizability to increase with the curation of larger golden call sets.

Another limitation of our approach relates to inconsistencies between srWGS read-depth signals and CNV labels derived from other sequencing technologies. In many challenging genomic regions, short-read coverage exhibits irregular and noisy fluctuations between CNV breakpoints defined by long-read or multi-technology callsets. These discrepancies often stem from mapping ambiguity, repetitive sequence structure, or short-read–specific biases. Consequently, although CHALLENGER reliably detects the presence of a CNV event, it struggles to uncover its precise boundaries within these problematic regions. To avoid penalizing the model with inconsistent boundary annotations, we removed the token level classification component during fine tuning. As future work, we aim to mitigate this limitation by incorporating additional representations such as paired-end mapping and split-read to improve breakpoint localization and report it alongside the gene level CNV class.

## 4 Methods

### 4.1 Datasets

#### Data Availability

The samples, ground truth data, and CNV predictions to reproduce the analyses can be accessed through https://zenodo.org/records/17593221.

#### Pre-Training sets from the samples in the 1000 Genomes Project

We pretrain the model using 550 randomly selected WGS samples from the 1000 Genomes Project. The sample IDs are listed in the Supplementary Table 24. CNV calls for these samples were obtained from the 1000 Genomes Phase 3 Reanalysis using DRAGEN v3.5, accessed in January 2025 from https://registry.opendata.aws/ilmn-dragen-1kgp. Training samples are labeled with 24,844 deletion (DEL) and 21,619 duplication (DUP) calls in total.

#### Fine tuning and test sets from the Schloissnig et al. call set

Schloissnig et al. [36] conducted a comprehensive analysis of structural variants (SVs) within the 1000 Genomes Project (1kGP) cohort using ONT (lrWGS). The study comprised 1,019 genomes sequenced to a median coverage of 16.9× with a median N50 read length of 20.3 kb. The authors introduced SAGA, a computational framework that integrates Sniffles v2.0.7 and an LRS-optimized implementation of DELLY v1.1.7 for SV discovery. In our work, we used 230 samples for fine-tuning and 30 samples for testing and benchmarking (Listed in Supplementary Tables 25 and 26). After filtering out CNVs smaller than 1 kbp, the training set contained 90,027 DEL and 12,162 DUP calls, while the test set included 11,990 DEL and 2,775 DUP calls.

#### Fine tuning and test sets from the Chaisson et al. call set

Chaisson et al. provide an integrated structural variant (SV) callset for 9 samples from 1000 Genomes Project (HG00512, HG00513, HG00731, HG00732, HG00733, NA19238, NA19239, NA19240) [32]. The callset includes CNVs detected using Illumina short-read WGS, 3.5 kb and 7.5 kb jumping libraries, PacBio long-read sequencing, and optical mapping. Additional technologies used for long-range phasing and haplotyping include 10X Chromium, Illumina synthetic long reads, Hi-C, and single-cell/single-strand sequencing. They use 15 WGS-based CNV callers for CNV detection. From the nine available samples, we select 6 (HG00513, HG00732, HG00733, NA19239, HG00512, HG00514) to fine-tune CHALLENGER, which includes 1,576 DEL and 3,066 DUP calls. We use the remaining 3 samples for inference and benchmarking, with a total of 786 DEL and 1,671 DUP events (See Supplementary Table 27).

#### Fine tuning and test sets in experimentally validated gene specific call sets

We fine-tune our base model using experimentally validated CNV call sets for four duplicated genes. For *SMN1/2*, we incorporate CNV events identified through Multiplex Ligation-dependent Probe Amplification (MLPA), resulting in a fine-tuning dataset of 20 samples (3 DEL, 5 DUP, and 12 with no event) and a test set of 21 samples (3 DEL, 6 DUP, and 12 with no event) for *SMN1*, and fine-tuning dataset of 24 samples (6 DEL, 6 DUP, and 12 with no event) and a test set of 24 samples (6 DEL, 6 DUP, and 12 with no event) for *SMN2*. To enable aggregate CNV prediction, we sum the read-depth signals of *SMN1* and *SMN2*. For *NPY4RA/B*, we use CNV labels derived from droplet digital PCR (ddPCR), comprising 9 samples for fine-tuning (5 DUP, and 4 with no event) and 9 samples for testing (6 DUP, and 3 with no event). For aggregate CNV fine-tuning, we merge the read-depth signals of *NPY4RA* and *NPY4RB*. As the original dataset does not include paralog-specific copy number annotations, it is excluded from the paralog-level CNV detection analysis. For *AMY1A/B/C* and *AMY2A*, we use CNV call sets generated from paralogue ratio tests (PRTs), duplication junction PCR, and microsatellite assays, yielding a fine-tuning dataset of 26 samples (7 DEL, 7 DUP, and 12 with no event) and a test set of 20 samples (5 DEL, 5 DUP, and 10 with no event) for *AMY1A/B/C*, and fine-tuning dataset of 24 samples (6 DEL, 6 DUP, and 12 with no event) and a test set of 22 samples (5 DEL, 6 DUP, and 11 with no event) for *AMY2A*. The original dataset provides aggregate copy numbers for *AMY1A/B/C* and paralog-specific copy numbers for *AMY2A*, and the analyses are conducted accordingly. All samples are listed in Supplementary Tables 28, 29, and 30.

### 4.2 Problem Formulation

Let *X* be the set of all genes with available read-depth (RD) signals, and let *X*^*i*^ denote the *i*^*th*^ gene where *i* ∈ {1, 2, …, *N* } and *N* = |*X*|. Each gene sample *X*^*i*^ is associated with two features (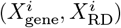), where 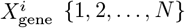 is the corresponding gene of *X*^*i*^ such that 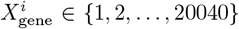 and 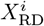 represents the standardized read-depth signal corresponding to that gene.

For standardization, the read-depth signals are first computed by averaging the sequencing coverage within 50 bp bins. For very short genomic regions, up to 15 kb of flanking sequence is appended to both upstream and downstream regions. Since the input vector size is fixed at 1000, genes longer than 50 kbp cannot fit into the vector when using 50 bp bins. For these cases, larger bin sizes are adaptively applied so that each gene’s read-depth signal fits within the 1000-length vector. Finally, each 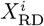 is left-padded with − 1 values to ensure a uniform input length of 1000 for all genes. For normalization, the average no-call signal across all genes is computed using 550 training samples, resulting in a look-up table for normalization. This search table, along with the pre-computed standard deviation value, is used to normalize the read-depth signal of each gene.

Let 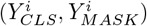 denote the ground-truth CNV label for *X*^*i*^ and the labels of the masked tokens in the same gene, respectively. The predictions of the model for *X*^*i*^ are given by 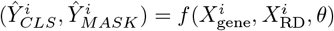. Here, *f* denotes a multi-class classifier parameterized by *θ*. The objective is to find the optimal parameters *θ*^∗^ that minimize the discrepancy between the predicted and ground-truth CNV labels in both gene-level and masked regions, i.e., 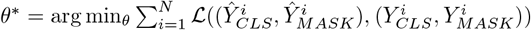, where ℒ denotes the loss function.

### 4.3 Model Description

The model first receives as input the normalized read-depth signal 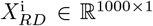 derived from a given gene region. This signal is transformed through two consecutive one-dimensional convolutional layers (1D CNNs). The first convolutional layer employs 32 filters (input channels = 1, output channels = 32), while the second layer employs 768 filters (input channels = 32, output channels = 768). Both layers use a kernel size of 3 and a stride of 1, and each is followed by batch normalization (BN) and a rectified linear unit (ReLU) activation. This operation produces an embedded representation *R*^*i*^ ∈ ℝ^1000*×*768^. A trainable classification token *E*_CLS_ ∈ ℝ^1*×*768^ and separator tokens *E*_SEP_ ∈ ℝ^1*×*768^ are then appended to the sequence, yielding a final read depth embedding of dimension ℝ^1002*×*768^.

To teach the model about the CNVs, the read depth embedding is element-wise summed with a corresponding CNV embedding representing one of four discrete states: *NOCALL, DEL, DUP*, or *MASK*. To enable gene-specific modeling of read-depth characteristics, the resulting tensor is further element-wise summed with a gene embedding vector 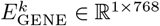 where 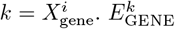 is a row in the learnable gene embedding matrix *E*_GENE_ ∈ ℝ^20040*×*768^ A positional embedding is subsequently added to preserve the sequential structure of the input signal. The resulting tensor is passed through a stack of three transformer encoder blocks, with an attention head count of 3, producing (i) the final hidden representation of the special [CLS] token, *C*^*i*^ ∈ ℝ^1*×*768^, and (ii) the final hidden representations of the read-depth tokens, *T* ^*i*^ ∈ ℝ^1000*×*768^.

CHALLENGER optimizes two complementary objectives through a Gene-Level Classification Head and Token-Level Mask Classification Head. The Classification Head processes the [CLS] representation *C*^*i*^ through a multilayer perceptron (MLP). The MLP layer first applies a LayerNorm, followed by a linear layer that keeps the hidden dimension at 768 units and a GELU activation function. A second linear layer then maps the 768-dimensional representation to 3 output units corresponding to the three CNV classes. A softmax over these three outputs produces probabilities for *DEL, DUP*, and *NO-CALL* states. The predicted CNV label for gene corresponds to the class with the highest probability, and this head is trained using a cross-entropy loss (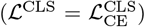). Token-Level Mask Classification Head operates on the final hidden representations of the read-depth tokens, *T* ^*i*^, passing it through a separate MLP to predict CNV labels for masked tokens. The MLP layer begins with a LayerNorm applied to each token representation, followed by a linear layer with 768 hidden units and a GELU activation function. A subsequent linear layer maps this transformed representation to three output units corresponding to the *DEL, DUP*, and *NO-CALL* classes. Applying a softmax to these outputs yields token-level CNV probabilities for each masked position. This head is trained using a combination of cross-entropy and Dice losses [41], where the Dice loss encourages spatial consistency and region-level overlap between predicted and ground-truth CNV segments (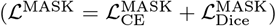). The final training objective is defined as the sum of the losses from both heads, ℒ_total_ = *λ*_cls_ ℒ^CLS^ + *λ*_mask_ ℒ^MASK^, where *λ*_*cls*_ and *λ*_*mask*_ are weighting coefficients controlling the relative contributions of the classification and segmentation objectives, respectively. We set *λ*_*cls*_ = 2.0, and *λ*_*mask*_ = 5.0.

### 4.4 Ablation Studies

We evaluated the contribution of CHALLENGER’s core components by removing the Transformer encoder, gene embeddings, and positional embeddings. All modified architectures are pre-trained and fine-tuned on the same datasets used for CHALLENGER^LR^ to ensure a fair comparison. Supplementary Table 23 summarizes the F1 performance of CHALLENGER^LR^ with these variant architectures on challenging genomic regions. Eliminating the Transformer encoder reduced the overall F1 score by 13%, highlighting the role of long-range contextual modeling. Removing gene embeddings caused the largest degradation with a 26% drop in overall F1 score, reflecting the critical importance of gene-level coverage characteristics, especially in challenging genomic regions. Excluding positional embeddings resulted in a 10% decrease in overall F1 score, indicating that preserving positional structure improves interpretation of read-depth patterns. Together, these results show that accurate CNV detection, particularly in complex, repetitive, and low-mappability genes, requires a multi-component architecture and demonstrate why CHALLENGER’s combination of convolutional layers, gene-aware embeddings, positional encodings, and transformer blocks is essential for robust performance.

### 4.5 Experimental Setup

#### Compared Methods

To evaluate CHALLENGER, we compare its performance against a comprehensive set of CNV callers spanning short-read, long-read, and paralog-specific analysis tools. For short-read WGS–based methods, we include DRAGEN [12], CNVnator [8], CUE [15], DELLY [28], LUMPY [17], and GATK [14]. DRAGEN is also used as the semi–ground truth source during pre-training. For long-read WGS–based methods, we benchmark against CuteSV [23], Severus [42], and SVision-Pro [25]. In experiments involving duplicated and paralogous genomic regions, we additionally compare CHALLENGER^GENE^ with tools specifically designed for paralog-aware CNV estimation, including Parascopy [35], QuicK-mer2 [40], and SMNCopyNumberCaller [37]. These tools are capable of aggregate or paralog-specific copy number inference. We run all compared tools with their default parameter settings.

#### Pre-Training

We pre-trained our model on 550 randomly selected whole-genome sequencing (WGS) samples from the 1000 Genomes Project for 1.5 epochs (∼400k steps), using DRAGEN-based CNV event calls. Challenging genes were excluded from the pre-training dataset. During the first 30k steps, 15% of CNVs were masked, followed by 30% masking for the next 30k steps, 60% for the next 30k steps, and 90% for the following 30k steps. The remaining steps were trained with full (100%) masking. We use Adam optimizer [43] and linear learning rate scheduler with an initial learning rate of 5*x*10^−5^.

#### Fine-Tuning

In fine-tuning, we fully mask the CNV labels across the entire regions of read depths and exclude the token-level classification head. This is motivated by the fact that CNV boundaries are often ambiguous due to noisy read-depth signals particularly in challenging regions. Penalizing the model with inconsistent CNV labels may prevent it from learning distinct CNV patterns. Therefore, we skip the token-level classification during fine-tuning. We use Adam optimizer [43] and a linear learning rate scheduler with an initial learning rate of 5 *×* 10^−5^ for all fine-tuning procedures. We always fine-tune the same pretrained model. ***CHALLENGER***^***LR***^: We fine-tune using 230 samples (300k steps). ***CHALLENGER***^***EXP***^: We fine-tuned with 6 samples (100k steps). ***CHALLENGER***^***GENE***^: *SMN1/2* with 44 samples (750 steps); *SMN1* with 20 samples (1200 steps); *SMN2* with 24 samples (400 steps); *AMY2A* with 24 samples (850 steps); *NPY4RA/B* with 9 samples (600 steps); *AMY1A/B/C* model with 26 samples (250 steps).

## References

1. Tuomo Mantere, Simone Kersten, and Alexander Hoischen. Long-read sequencing emerging in medical genetics. Frontiers in genetics, 10:432668, 2019.

2. Birgitt Schüle, Karen N McFarland, Kelsey Lee, Yu-Chih Tsai, Khanh-Dung Nguyen, Chao Sun, Mei Liu, Christie Byrne, Ramesh Gopi, Neng Huang, et al. Parkinson’s disease associated with pure atxn10 repeat expansion. NPJ Parkinson’s disease, 3(1):27, 2017.

3. Karen N McFarland, Jilin Liu, Ivette Landrian, Ronald Godiska, Savita Shanker, Fahong Yu, William G Farmerie, and Tetsuo Ashizawa. Smrt sequencing of long tandem nucleotide repeats in sca10 reveals unique insight of repeat expansion structure. PloS one, 10(8):e0135906, 2015.

4. O Pös, J Radvanszky, G Buglyó, Z Pös, D Rusnakova, B Nagy, and T Szemes. Dna copy number variation: main characteristics, evolutionary significance, and pathological aspects. biomed j 44: 548–559, 2021.

5. Rebecca Truty, Joshua Paul, Michael Kennemer, Stephen E Lincoln, Eric Olivares, Robert L Nussbaum, and Swaroop Aradhya. Prevalence and properties of intragenic copy-number variation in mendelian disease genes. Genetics in Medicine, 21(1):114–123, 2019.

6. Quentin Testard, Xavier Vanhoye, Kevin Yauy, Marie-Emmanuelle Naud, Gaelle Vieville, Francis Rousseau, Benjamin Dauriat, Valentine Marquet, Sylvie Bourthoumieu, David Geneviève, et al. Exome sequencing as a first-tier test for copy number variant detection: retrospective evaluation and prospective screening in 2418 cases. Journal of medical genetics, 59(12):1234–1240, 2022.

7. Johannes Smolander, Sofia Khan, Kalaimathy Singaravelu, Leni Kauko, Riikka J Lund, Asta Laiho, and Laura L Elo. Evaluation of tools for identifying large copy number variations from ultra-low-coverage whole-genome sequencing data. BMC Genomics, 22(1):357, May 2021.

8. Alexej Abyzov, Alexander E Urban, Michael Snyder, and Mark Gerstein. Cnvnator: an approach to discover, genotype, and characterize typical and atypical cnvs from family and population genome sequencing. Genome research, 21(6):974–984, 2011.

9. Valentina Boeva, Tatiana Popova, Kevin Bleakley, Pierre Chiche, Julie Cappo, Gudrun Schleiermacher, Isabelle Janoueix-Lerosey, Olivier Delattre, and Emmanuel Barillot. Control-FREEC: a tool for assessing copy number and allelic content using next-generation sequencing data. Bioinformatics, 28(3):423–425, 12 2011.

10. Berk Mandiracioglu, Furkan Ozden, Gun Kaynar, Mehmet Alper Yilmaz, Can Alkan, and A Ercument Cicek. Ecole: Learning to call copy number variants on whole exome sequencing data. Nature Communications, 15(1):132, 2024.

11. Ramesh Rajaby and Wing-Kin Sung. Survindel2: improving copy number variant calling from next-generation sequencing using hidden split reads. Nature Communications, 15(1):1–16, 2024.

12. Sairam Behera, Severine Catreux, Massimiliano Rossi, Sean Truong, Zhuoyi Huang, Michael Ruehle, Arun Visvanath, Gavin Parnaby, Cooper Roddey, Vitor Onuchic, et al. Comprehensive genome analysis and variant detection at scale using dragen. Nature Biotechnology, 43(7):1177–1191, 2025.

13. Furkan Özden, Can Alkan, and A Ercüment Çiçek. Polishing copy number variant calls on exome sequencing data via deep learning. Genome research, 32(6):1170–1182, 2022.

14. Aaron McKenna, Matthew Hanna, Eric Banks, Andrey Sivachenko, Kristian Cibulskis, Andrew Kernytsky, Kiran Garimella, David Altshuler, Stacey Gabriel, Mark Daly, et al. The genome analysis toolkit: a mapreduce framework for analyzing next-generation dna sequencing data. Genome research, 20(9):1297–1303, 2010.

15. Victoria Popic, Chris Rohlicek, Fabio Cunial, Iman Hajirasouliha, Dmitry Meleshko, Kiran Garimella, and Anant Maheshwari. Cue: a deep-learning framework for structural variant discovery and genotyping. Nature Methods, 20(4):559–568, 2023.

16. Samantha Zarate, Andrew Carroll, Medhat Mahmoud, Olga Krasheninina, Goo Jun, William J Salerno, Michael C Schatz, Eric Boerwinkle, Richard A Gibbs, and Fritz J Sedlazeck. Parliament2: Accurate structural variant calling at scale. GigaScience, 9(12):giaa145, 2020.

17. Ryan M Layer, Colby Chiang, Aaron R Quinlan, and Ira M Hall. Lumpy: a probabilistic framework for structural variant discovery. Genome biology, 15(6):R84, 2014.

18. Mark TW Ebbert, Tanner D Jensen, Karen Jansen-West, Jonathon P Sens, Joseph S Reddy, Perry G Ridge, John SK Kauwe, Veronique Belzil, Luc Pregent, Minerva M Carrasquillo, et al. Systematic analysis of dark and camouflaged genes reveals disease-relevant genes hiding in plain sight. Genome biology, 20(1):97, 2019.

19. Francesco Kumara Mastrorosa, Danny E Miller, and Evan E Eichler. Applications of long-read sequencing to mendelian genetics. Genome Medicine, 15(1):42, 2023.

20. Josephine B Oehler, Helen Wright, Zornitza Stark, Andrew J Mallett, and Ulf Schmitz. The application of long-read sequencing in clinical settings. Human genomics, 17(1):73, 2023.

21. Susan M Hiatt, James MJ Lawlor, Lori H Handley, Ryne C Ramaker, Brianne B Rogers, E Christopher Partridge, Lori Beth Boston, Melissa Williams, Christopher B Plott, Jerry Jenkins, et al. Long-read genome sequencing for the molecular diagnosis of neurodevelopmental disorders. Human Genetics and Genomics Advances, 2(2), 2021.

22. Fritz J Sedlazeck, Philipp Rescheneder, Moritz Smolka, Han Fang, Maria Nattestad, Arndt Von Haeseler, and Michael C Schatz. Accurate detection of complex structural variations using single-molecule sequencing. Nature methods, 15(6):461–468, 2018.

23. Tao Jiang, Shiqi Liu, Shuqi Cao, and Yadong Wang. Structural variant detection from long-read sequencing data with cutesv. In Variant Calling: Methods and Protocols, pages 137–151. Springer, 2022.

24. Luca Denti, Parsoa Khorsand, Paola Bonizzoni, Fereydoun Hormozdiari, and Rayan Chikhi. Svdss: structural variation discovery in hard-to-call genomic regions using sample-specific strings from accurate long reads. Nature Methods, 20(4):550–558, 2023.

25. Jiadong Lin, Songbo Wang, Peter A Audano, Deyu Meng, Jacob I Flores, Walter Kosters, Xiaofei Yang, Peng Jia, Tobias Marschall, Christine R Beck, et al. Svision: a deep learning approach to resolve complex structural variants. Nature methods, 19(10):1230–1233, 2022.

26. Yu Chen, Amy Y Wang, Courtney A Barkley, Yixin Zhang, Xinyang Zhao, Min Gao, Mick D Edmonds, and Zechen Chong. Deciphering the exact breakpoints of structural variations using long sequencing reads with debreak. Nature Communications, 14(1):283, 2023.

27. Christopher T Saunders, James M Holt, Daniel N Baker, Juniper A Lake, Jonathan R Belyeu, Zev Kronenberg, William J Rowell, and Michael A Eberle. Sawfish: Improving long-read structural variant discovery and genotyping with local haplotype modeling. Bioinformatics, 41(4):btaf136, 2025.

28. Tobias Rausch, Thomas Zichner, Andreas Schlattl, Adrian M Stütz, Vladimir Benes, and Jan O Korbel. Delly: structural variant discovery by integrated paired-end and split-read analysis. Bioinformatics, 28(18):i333–i339, 2012.

29. Patrick Marks, Sarah Garcia, Alvaro Martinez Barrio, Kamila Belhocine, Jorge Bernate, Rajiv Bharadwaj, Keith Bjornson, Claudia Catalanotti, Josh Delaney, Adrian Fehr, et al. Resolving the full spectrum of human genome variation using linked-reads. Genome research, 29(4):635–645, 2019.

30. Xuefang Zhao, Ryan L Collins, Wan-Ping Lee, Alexandra M Weber, Yukyung Jun, Qihui Zhu, Ben Weisburd, Yongqing Huang, Peter A Audano, Harold Wang, et al. Expectations and blind spots for structural variation detection from long-read assemblies and short-read genome sequencing technologies. The American Journal of Human Genetics, 108(5):919–928, 2021.

31. Mehmet Alper Yılmaz, Ahmet Arda Ceylan, Gün Kaynar, and A Ercüment Çiçek. Lyceum: Learning to call copy number variants on low coverage ancient genomes. bioRxiv, pages 2024–10, 2024.

32. Mark JP Chaisson, Ashley D Sanders, Xuefang Zhao, Ankit Malhotra, David Porubsky, Tobias Rausch, Eugene J Gardner, Oscar L Rodriguez, Li Guo, Ryan L Collins, et al. Multi-platform discovery of haplotype-resolved structural variation in human genomes. Nature communications, 10(1):1784, 2019.

33. Marta Byrska-Bishop, Uday S Evani, Xuefang Zhao, Anna O Basile, Haley J Abel, Allison A Regier, André Corvelo Wayne E Clarke, Rajeeva Musunuri, Kshithija Nagulapalli, et al. High-coverage whole-genome sequencing of the expanded 1000 genomes project cohort including 602 trios. Cell, 185(18):3426–3440, 2022.

34. Yinhan Liu, Myle Ott, Naman Goyal, Jingfei Du, Mandar Joshi, Danqi Chen, Omer Levy, Mike Lewis, Luke Zettlemoyer, and Veselin Stoyanov. Roberta: A robustly optimized bert pretraining approach. arXiv preprint 1907.11692, 2019.

35. Timofey Prodanov and Vikas Bansal. Robust and accurate estimation of paralog-specific copy number for duplicated genes using whole-genome sequencing. Nature Communications, 13(1):3221, 2022.

36. Siegfried Schloissnig, Samarendra Pani, Jana Ebler, Carsten Hain, Vasiliki Tsapalou, Arda Söylev, Patrick Hüther, Hufsah Ashraf, Timofey Prodanov, Mila Asparuhova, et al. Structural variation in 1,019 diverse humans based on long-read sequencing. Nature, pages 1–11, 2025.

37. Xiao Chen, Alba Sanchis-Juan, Courtney E French, Andrew J Connell, Isabelle Delon, Zoya Kingsbury, Aditi Chawla, Aaron L Halpern, Ryan J Taft, NIHR BioResource, et al. Spinal muscular atrophy diagnosis and carrier screening from genome sequencing data. Genetics in Medicine, 22(5):945–953, 2020.

38. Kateryna Shebanits, Torsten Günther, Anna CV Johansson, Khurram Maqbool, Lars Feuk, Mattias Jakobsson, and Dan Larhammar. Copy number determination of the gene for the human pancreatic polypeptide receptor npy4r using read depth analysis and droplet digital pcr. BMC biotechnology, 19(1):31, 2019.

39. Danielle Carpenter, Sugandha Dhar, Laura M Mitchell, Beiyuan Fu, Jess Tyson, Nzar AA Shwan, Fengtang Yang, Mark G Thomas, and John AL Armour. Obesity, starch digestion and amylase: association between copy number variants at human salivary (amy1) and pancreatic (amy2) amylase genes. Human molecular genetics, 24(12):3472–3480, 2015.

40. F Shen and JM Kidd. Rapid, paralog-sensitive cnv analysis of 2457 human genomes using quick-mer2. genes (basel). 2020: 11 (2): 141.

41. Fausto Milletari, Nassir Navab, and Seyed-Ahmad Ahmadi. V-net: Fully convolutional neural networks for volumetric medical image segmentation. In 2016 fourth international conference on 3D vision (3DV), pages 565–571. Ieee, 2016.

42. Ayse Keskus, Asher Bryant, Tanveer Ahmad, Byunggil Yoo, Sergey Aganezov, Anton Goretsky, Ataberk Donmez, Lisa A Lansdon, Isabel Rodriguez, Jimin Park, et al. Severus: accurate detection and characterization of somatic structural variation in tumor genomes using long reads. medRxiv, 2024.

43. Diederik P Kingma. Adam: A method for stochastic optimization. arXiv preprint 1412.6980, 2014.

