## Supplementary Material for "CHALLENGER: Detecting Copy Number Variants in Challenging Regions Using Whole Genome Sequencing Data"

### Supplementary Material
### for
### CHALLENGER: Detecting Copy Number Variants in Challenging Genes Using Whole Genome Sequencing Data

#### 1 Supplementary Figures

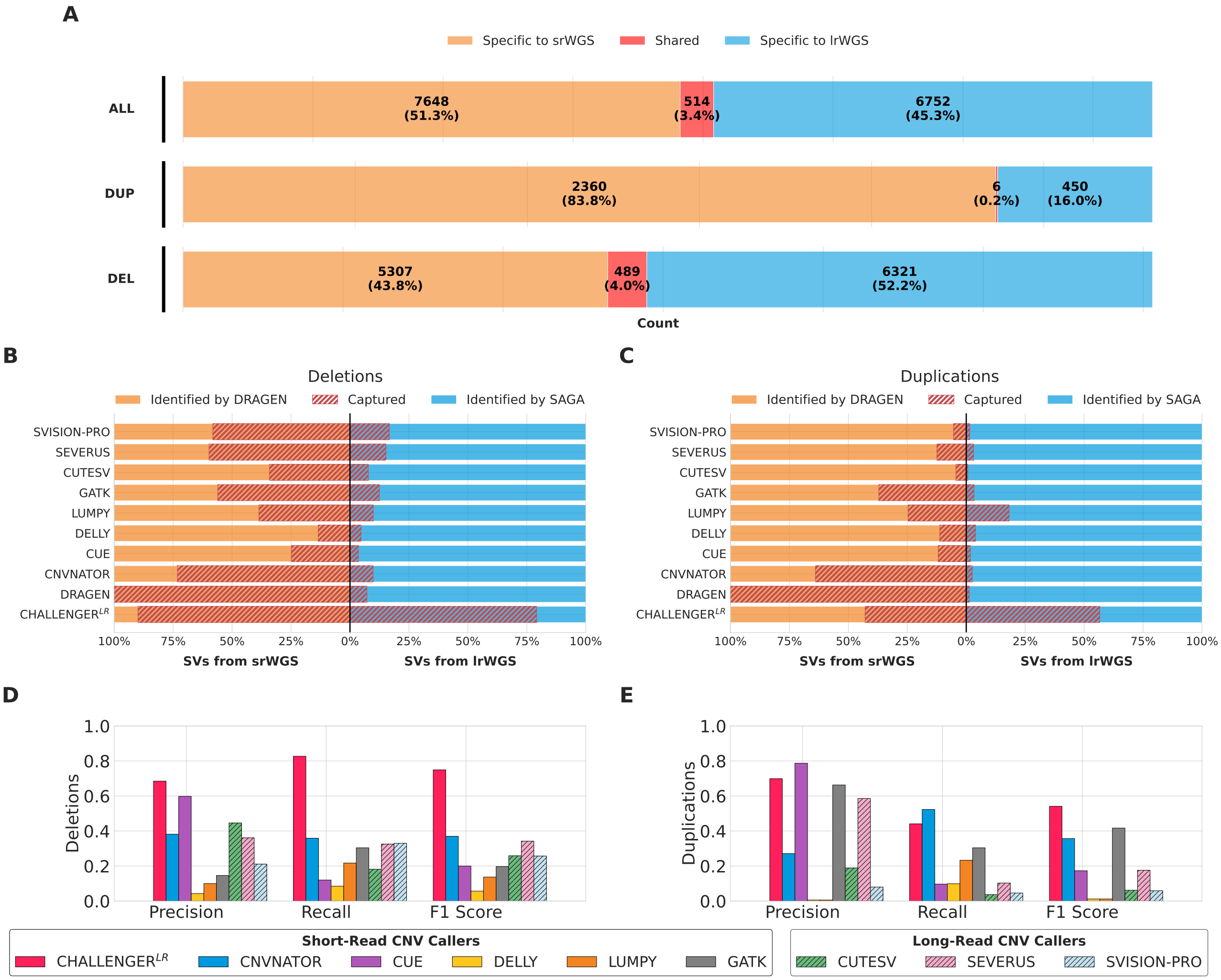


**Supplementary Figure 1.** Evaluation of CHALLENGER$^{\mathrm{LR}}$ in detecting CNVs specific to long-read and short-read sequencing within all genomic regions. **(A)** Overlap between CNV calls derived from short-read WGS (srWGS; DRAGEN) and long-read WGS (lrWGS; SAGA) across all genes. **(B–C)** Comparative recall of CHALLENGER$^{\mathrm{LR}}$ and existing methods for CNVs identified in srWGS and lrWGS datasets, shown separately for deletions (B) and duplications (C). **(D–E)** Comparison of precision, recall, and F1-score across methods for deletions (D) and duplications (E) within all gene regions.

#### 2 Supplementary Tables

**Supplementary Table 1.** Recall of deletion (DEL) and duplication (DUP) events in challenging genomic regions using short-read CNV calls from DRAGEN and long-read CNV calls from SAGA.

| **TOOLS** | **Short-Read DEL Recall** | **Long-Read DEL Recall** | **Short-Read DUP Recall** | **Long-Read DUP Recall** |
|---|---|---|---|---|
| CHALLENGER$^{LR}$ | 0.882 | **0.833** | 0.682 | **0.258** |
| DRAGEN | **1.000** | 0.120 | **1.000** | 0.021 |
| CNVNATOR | 0.674 | 0.172 | 0.616 | 0.052 |
| CUE | 0.230 | 0.072 | 0.147 | 0.041 |
| DELLY | 0.114 | 0.040 | 0.078 | 0.052 |
| LUMPY | 0.370 | 0.136 | 0.181 | 0.144 |
| GATK | 0.516 | 0.231 | 0.231 | 0.082 |
| CUTESV | 0.332 | 0.139 | 0.072 | 0.010 |
| SEVERUS | 0.554 | 0.264 | 0.160 | 0.072 |
| SVISION-PRO | 0.563 | 0.276 | 0.071 | 0.021 |

**Supplementary Table 2.** Performance comparison of WGS-based CNV callers on the 1000 Genomes test set within gene regions located in challenging genomic regions. The union of DRAGEN calls from short-read WGS (srWGS) and SAGA calls from long-read WGS (lrWGS) is used as the ground truth. Bold values indicate the best performance in each metric.

| **TOOLS** | **DEL Prec** | **DUP Prec** | **Overall Prec** | **DEL Rec** | **DUP Rec** | **Overall Rec** | **DEL F1** | **DUP F1** | **Overall F1** |
|---|---|---|---|---|---|---|---|---|---|
| CNVNATOR | 0.315 | 0.149 | 0.232 | 0.396 | 0.527 | 0.462 | 0.351 | 0.232 | 0.291 |
| CUE | 0.494 | **0.719** | 0.607 | 0.134 | 0.129 | 0.131 | 0.211 | 0.219 | 0.215 |
| DELLY | 0.118 | 0.011 | 0.064 | 0.079 | 0.072 | 0.075 | 0.094 | 0.019 | 0.057 |
| LUMPY | 0.221 | 0.010 | 0.115 | 0.243 | 0.177 | 0.210 | 0.232 | 0.018 | 0.125 |
| GATK | 0.113 | 0.645 | 0.379 | 0.354 | 0.207 | 0.280 | 0.171 | 0.313 | 0.242 |
| CUTESV | 0.416 | 0.175 | 0.295 | 0.218 | 0.062 | 0.140 | 0.286 | 0.092 | 0.189 |
| SEVERUS | 0.356 | 0.577 | 0.466 | 0.375 | 0.146 | 0.260 | 0.365 | 0.233 | 0.299 |
| SVISION-PRO | 0.277 | 0.140 | 0.209 | 0.392 | 0.066 | 0.229 | 0.325 | 0.090 | 0.207 |
| CHALLENGER$^{LR}$ | **0.725** | 0.650 | **0.688** | **0.844** | **0.618** | **0.731** | **0.780** | **0.634** | **0.707** |

**Supplementary Table 3.** Confusion matrices for the performance comparison of WGS-based CNV callers on the 1000 Genomes test set within gene regions located in challenging genomic regions. The union of DRAGEN calls from short-read WGS (srWGS) and SAGA calls from long-read WGS (lrWGS) is used as the ground truth. These produce the precision and recall results presented in Supplementary Table 2.

| | | **Ground Truth** | | |
|---|---|---|---|---|
| TOOLS | **Predicted** | NO CALL | DUP | DEL |
| CNVNATOR | NO CALL | 90491 | 344 | 3369 |
| | DUP | 2303 | 408 | 35 |
| | DEL | 4836 | 22 | 2232 |
| CUE | NO CALL | 96843 | 651 | 4882 |
| | DUP | 39 | 100 | 0 |
| | DEL | 748 | 23 | 754 |
| DELLY | NO CALL | 89374 | 704 | 5067 |
| | DUP | 4962 | 56 | 126 |
| | DEL | 3294 | 14 | 443 |
| LUMPY | NO CALL | 79314 | 619 | 3784 |
| | DUP | 13506 | 137 | 482 |
| | DEL | 4810 | 18 | 1370 |
| GATK | NO CALL | 81934 | 532 | 3639 |
| | DUP | 84 | 160 | 4 |
| | DEL | 15612 | 82 | 1993 |
| CUTESV | NO CALL | 95765 | 647 | 4398 |
| | DUP | 217 | 48 | 9 |
| | DEL | 1648 | 79 | 1229 |
| SEVERUS | NO CALL | 93818 | 564 | 3523 |
| | DUP | 82 | 113 | 1 |
| | DEL | 3730 | 97 | 2112 |
| SVISION-PRO | NO CALL | 91730 | 573 | 3413 |
| | DUP | 299 | 51 | 15 |
| | DEL | 5601 | 150 | 2208 |
| CHALLENGER$^{LR}$ | NO CALL | 95579 | 287 | 853 |
| | DUP | 228 | 478 | 29 |
| | DEL | 1793 | 9 | 4754 |

**Supplementary Table 4.** Recall of deletion (DEL) and duplication (DUP) events in all genomic regions using short-read CNV calls from DRAGEN and long-read CNV calls from SAGA.

| **TOOLS** | **Short-Read DEL Recall** | **Long-Read DEL Recall** | **Short-Read DUP Recall** | **Long-Read DUP Recall** |
|---|---|---|---|---|
| CHALLENGER$^{LR}$ | 0.899 | **0.792** | 0.429 | **0.566** |
| DRAGEN | **1.000** | 0.072 | **1.000** | 0.013 |
| CNVNATOR | 0.732 | 0.098 | 0.639 | 0.026 |
| CUE | 0.248 | 0.036 | 0.119 | 0.018 |
| DELLY | 0.134 | 0.047 | 0.112 | 0.039 |
| LUMPY | 0.386 | 0.099 | 0.246 | 0.182 |
| GATK | 0.561 | 0.125 | 0.371 | 0.033 |
| CUTESV | 0.341 | 0.078 | 0.044 | 0.007 |
| SEVERUS | 0.598 | 0.152 | 0.124 | 0.031 |
| SVISION-PRO | 0.582 | 0.167 | 0.054 | 0.015 |

**Supplementary Table 5.** Performance comparison of WGS-based CNV callers on the 1000 Genomes test set within gene regions located in all genomic regions. The union of DRAGEN calls from short-read WGS (srWGS) and SAGA calls from long-read WGS (lrWGS) is used as the ground truth. Bold values indicate the best performance in each metric.

| TOOLS | DEL Prec | DUP Prec | Overall Prec | DEL Rec | DUP Rec | Overall Rec | DEL F1 | DUP F1 | Overall F1 |
|---|---|---|---|---|---|---|---|---|---|
| CNVNATOR | 0.382 | 0.271 | 0.327 | 0.359 | **0.523** | 0.441 | 0.370 | 0.357 | 0.364 |
| CUE | 0.598 | **0.787** | **0.692** | 0.120 | 0.097 | 0.109 | 0.200 | 0.173 | 0.186 |
| DELLY | 0.043 | 0.006 | 0.025 | 0.085 | 0.099 | 0.092 | 0.057 | 0.012 | 0.034 |
| LUMPY | 0.100 | 0.006 | 0.053 | 0.217 | 0.233 | 0.225 | 0.137 | 0.012 | 0.074 |
| GATK | 0.146 | 0.663 | 0.404 | 0.304 | 0.304 | 0.304 | 0.197 | 0.417 | 0.307 |
| CUTESV | 0.446 | 0.189 | 0.318 | 0.182 | 0.037 | 0.109 | 0.259 | 0.062 | 0.160 |
| SEVERUS | 0.361 | 0.586 | 0.473 | 0.325 | 0.103 | 0.214 | 0.342 | 0.176 | 0.259 |
| SVISION-PRO | 0.211 | 0.080 | 0.145 | 0.330 | 0.046 | 0.188 | 0.257 | 0.059 | 0.158 |
| CHALLENGER$^{LR}$ | **0.685** | 0.699 | 0.692 | **0.827** | 0.441 | **0.634** | **0.749** | **0.541** | **0.645** |

**Supplementary Table 6.** Confusion matrices for the performance comparison of WGS-based CNV callers on the 1000 Genomes test set within gene regions located in all genomic regions. The union of DRAGEN calls from short-read WGS (srWGS) and SAGA calls from long-read WGS (lrWGS) is used as the ground truth. These produce the precision and recall results presented in Supplementary Table 5.

| | | **Ground Truth** | | |
|---|---|---|---|---|
| TOOLS | **Predicted** | NO CALL | DUP | DEL |
| CNVNATOR | NO CALL | 547789 | 1294 | 7645 |
| | DUP | 3866 | 1451 | 36 |
| | DEL | 6940 | 30 | 4309 |
| CUE | NO CALL | 557597 | 2464 | 10549 |
| | DUP | 73 | 269 | 0 |
| | DEL | 925 | 42 | 1441 |
| DELLY | NO CALL | 492463 | 2408 | 10509 |
| | DUP | 43413 | 274 | 458 |
| | DEL | 22719 | 93 | 1023 |
| LUMPY | NO CALL | 431067 | 2063 | 8144 |
| | DUP | 104166 | 646 | 1245 |
| | DEL | 23362 | 66 | 2601 |
| GATK | NO CALL | 536912 | 1838 | 8339 |
| | DUP | 424 | 843 | 5 |
| | DEL | 21259 | 94 | 3646 |
| CUTESV | NO CALL | 555551 | 2574 | 9795 |
| | DUP | 427 | 102 | 10 |
| | DEL | 2617 | 99 | 2185 |
| SEVERUS | NO CALL | 551646 | 2337 | 8083 |
| | DUP | 198 | 287 | 5 |
| | DEL | 6751 | 151 | 3902 |
| SVISION-PRO | NO CALL | 542526 | 2423 | 7997 |
| | DUP | 1454 | 129 | 36 |
| | DEL | 14615 | 223 | 3957 |
| CHALLENGER$^{LR}$ | NO CALL | 553543 | 1529 | 2023 |
| | DUP | 476 | 1224 | 51 |
| | DEL | 4546 | 22 | 9916 |

**Supplementary Table 7.** Performance comparison of WGS-based CNV callers on the Chaisson et al.'s human expert-curated calls within gene regions located in challenging genomic regions. Bold values indicate the best performance in each metric.

| **TOOLS** | **DEL Prec** | **DUP Prec** | **Overall Prec** | **DEL Rec** | **DUP Rec** | **Overall Rec** | **DEL F1** | **DUP F1** | **Overall F1** |
|---|---|---|---|---|---|---|---|---|---|
| DRAGEN | 0.550 | 0.200 | 0.375 | 0.200 | 0.032 | 0.116 | 0.294 | 0.055 | 0.175 |
| CNVNATOR | 0.392 | 0.194 | 0.293 | 0.315 | 0.149 | 0.232 | 0.349 | 0.169 | 0.259 |
| CUE | 0.642 | 0.154 | 0.398 | 0.119 | 0.005 | 0.062 | 0.200 | 0.010 | 0.105 |
| DELLY | 0.225 | 0.030 | 0.127 | 0.127 | 0.032 | 0.080 | 0.162 | 0.031 | 0.097 |
| LUMPY | 0.377 | 0.037 | 0.207 | 0.295 | 0.155 | 0.225 | 0.331 | 0.059 | 0.195 |
| GATK | 0.120 | 0.143 | 0.131 | 0.390 | 0.011 | 0.200 | 0.183 | 0.020 | 0.102 |
| CUTESV | 0.762 | 0.077 | 0.420 | 0.208 | 0.005 | 0.107 | 0.327 | 0.010 | 0.168 |
| SEVERUS | 0.662 | 0.102 | 0.382 | 0.472 | 0.016 | 0.244 | 0.551 | 0.028 | 0.289 |
| SVISION-PRO | 0.687 | 0.058 | 0.372 | 0.434 | 0.019 | 0.226 | 0.532 | 0.028 | 0.280 |
| CHALLENGER$^{EXP}$ | **0.844** | **0.879** | **0.861** | **0.754** | **0.869** | **0.812** | **0.797** | **0.874** | **0.835** |

**Supplementary Table 8.** Confusion matrices for the performance comparison of WGS-based CNV callers on the Chaisson et al.'s human expert-curated calls within gene regions located in challenging genomic regions. These produce the precision and recall results presented in Supplementary Table 7.

| | | **Ground Truth** | | |
|---|---|---|---|---|
| TOOLS | **Predicted** | NO CALL | DUP | DEL |
| DRAGEN | NO CALL | 9043 | 353 | 650 |
| | DUP | 44 | 12 | 4 |
| | DEL | 124 | 10 | 164 |
| CNVNATOR | NO CALL | 8609 | 292 | 556 |
| | DUP | 228 | 56 | 4 |
| | DEL | 374 | 27 | 258 |
| CUE | NO CALL | 9152 | 367 | 721 |
| | DUP | 11 | 2 | 0 |
| | DEL | 48 | 6 | 97 |
| DELLY | NO CALL | 8497 | 344 | 703 |
| | DUP | 374 | 12 | 11 |
| | DEL | 340 | 19 | 104 |
| LUMPY | NO CALL | 7361 | 301 | 524 |
| | DUP | 1468 | 58 | 53 |
| | DEL | 382 | 16 | 241 |
| GATK | NO CALL | 7025 | 193 | 497 |
| | DUP | 22 | 4 | 2 |
| | DEL | 2164 | 178 | 319 |
| CUTESV | NO CALL | 9142 | 367 | 646 |
| | DUP | 22 | 2 | 2 |
| | DEL | 47 | 6 | 170 |
| SEVERUS | NO CALL | 8982 | 349 | 431 |
| | DUP | 52 | 6 | 1 |
| | DEL | 177 | 20 | 386 |
| SVISION-PRO | NO CALL | 8964 | 346 | 457 |
| | DUP | 107 | 7 | 6 |
| | DEL | 140 | 22 | 355 |
| CHALLENGER$^{EXP}$ | NO CALL | 9072 | 42 | 185 |
| | DUP | 29 | 326 | 16 |
| | DEL | 107 | 7 | 617 |

**Supplementary Table 9.** Performance comparison of WGS-based CNV callers on the Chaisson et al.'s human expert-curated calls within gene regions located in all genomic regions. Bold values indicate the best performance in each metric.

| TOOLS | DEL Prec | DUP Prec | Overall Prec | DEL Rec | DUP Rec | Overall Rec | DEL F1 | DUP F1 | Overall F1 |
|---|---|---|---|---|---|---|---|---|---|
| DRAGEN | 0.618 | 0.225 | 0.421 | 0.206 | 0.043 | 0.125 | 0.309 | 0.073 | 0.191 |
| CNVNATOR | 0.453 | 0.173 | 0.313 | 0.289 | 0.115 | 0.202 | 0.353 | 0.138 | 0.245 |
| CUE | 0.672 | 0.215 | 0.444 | 0.103 | 0.018 | 0.060 | 0.179 | 0.033 | 0.106 |
| DELLY | 0.088 | 0.011 | 0.050 | 0.147 | 0.042 | 0.094 | 0.110 | 0.017 | 0.064 |
| LUMPY | 0.172 | 0.014 | 0.093 | 0.273 | 0.211 | 0.242 | 0.211 | 0.026 | 0.119 |
| GATK | 0.134 | 0.134 | 0.134 | 0.319 | 0.022 | 0.170 | 0.188 | 0.037 | 0.113 |
| CUTESV | 0.790 | 0.067 | 0.428 | 0.176 | 0.005 | 0.091 | 0.288 | 0.009 | 0.149 |
| SEVERUS | 0.731 | 0.032 | 0.381 | 0.442 | 0.024 | 0.233 | 0.551 | 0.028 | 0.289 |
| SVISION-PRO | 0.740 | 0.044 | 0.392 | 0.394 | 0.015 | 0.205 | 0.515 | 0.023 | 0.269 |
| CHALLENGER$^{EXP}$ | **0.802** | **0.889** | **0.846** | **0.749** | **0.874** | **0.811** | **0.775** | **0.881** | **0.828** |

**Supplementary Table 10.** Confusion matrices for the performance comparison of WGS-based CNV callers on the Chaisson et al.'s human expert-curated calls within gene regions located in all genomic regions. These produce the precision and recall results presented in Supplementary Table 9.

| | | **Ground Truth** | | |
|---|---|---|---|---|
| TOOLS | **Predicted** | NO CALL | DUP | DEL |
| DRAGEN | NO CALL | 54571 | 736 | 1321 |
| | DUP | 111 | 34 | 6 |
| | DEL | 197 | 16 | 344 |
| CNVNATOR | NO CALL | 53914 | 655 | 1182 |
| | DUP | 423 | 90 | 6 |
| | DEL | 542 | 41 | 483 |
| CUE | NO CALL | 54753 | 763 | 1499 |
| | DUP | 51 | 14 | 0 |
| | DEL | 75 | 9 | 172 |
| DELLY | NO CALL | 49461 | 714 | 1403 |
| | DUP | 2930 | 33 | 23 |
| | DEL | 2488 | 39 | 245 |
| LUMPY | NO CALL | 41055 | 587 | 1060 |
| | DUP | 11666 | 166 | 155 |
| | DEL | 2158 | 33 | 456 |
| GATK | NO CALL | 51536 | 548 | 1134 |
| | DUP | 106 | 17 | 4 |
| | DEL | 3237 | 221 | 533 |
| CUTESV | NO CALL | 54755 | 774 | 1375 |
| | DUP | 54 | 4 | 2 |
| | DEL | 70 | 8 | 294 |
| SEVERUS | NO CALL | 54067 | 740 | 929 |
| | DUP | 567 | 19 | 4 |
| | DEL | 245 | 27 | 738 |
| SVISION-PRO | NO CALL | 54428 | 748 | 998 |
| | DUP | 246 | 12 | 14 |
| | DEL | 205 | 26 | 659 |
| CHALLENGER$^{EXP}$ | NO CALL | 54519 | 85 | 397 |
| | DUP | 63 | 687 | 23 |
| | DEL | 294 | 14 | 1251 |

**Supplementary Table 11.** Performance comparison of duplicated-region copy number callers on ddPCR-validated NPY4R samples. Bold values highlight the top performer for each metric.

| GENE | TOOLS | DEL Prec | DUP Prec | Overall Prec | DEL Rec | DUP Rec | Overall Rec | DEL F1 | DUP F1 | Overall F1 |
|---|---|---|---|---|---|---|---|---|---|---|
| NPY4R/2 | PARASCOPY | - | **1.000** | - | - | 0.667 | - | - | 0.800 | - |
| | QUICKMER2 | - | **1.000** | - | - | 0.667 | - | - | 0.800 | - |
| | CHALLENGER$^{GENE}$ | **-** | **1.000** | - | **-** | **0.833** | - | **-** | **0.909** | **-** |

**Supplementary Table 12.** Confusion matrices for the performance comparison of duplicated-region copy number callers on ddPCR-validated NPY4R samples. These matrices correspond to the precision and recall values presented in 11

| | | | **Ground Truth** | | |
|---|---|---|---|---|---|
| **GENE** | **TOOL** | **Predicted** | NO CALL | DUP | DEL |
| NPY4R/2 | PARASCOPY | NO CALL | 3 | 2 | 0 |
| | | DUP | 0 | 4 | 0 |
| | | DEL | 0 | 0 | 0 |
| | QUICKMER2 | NO CALL | 3 | 2 | 0 |
| | | DUP | 0 | 4 | 0 |
| | | DEL | 0 | 0 | 0 |
| | CHALLENGER$^{GENE}$ | NO CALL | 3 | 1 | 0 |
| | | DUP | 0 | 5 | 0 |
| | | DEL | 0 | 0 | 0 |

**Supplementary Table 13.** Performance comparison of duplicated-region copy number callers on experimentally validated AMY1A/B/C samples. Bold values indicate the best-performing method for each metric.

| GENE | TOOLS | DEL Prec | DUP Prec | Overall Prec | DEL Rec | DUP Rec | Overall Rec | DEL F1 | DUP F1 | Overall F1 |
|---|---|---|---|---|---|---|---|---|---|---|
| AMY1 A/B/C | PARASCOPY | 0.833 | **1.000** | 0.917 | **1.000** | **0.800** | **0.900** | 0.909 | **0.889** | 0.899 |
| | QUICKMER2 | 0.357 | **1.000** | 0.679 | **1.000** | **0.800** | **0.900** | 0.526 | **0.889** | 0.708 |
| | CHALLENGER$^{GENE}$ | **1.000** | **1.000** | **1.000** | **1.000** | **0.800** | **0.900** | **1.000** | **0.889** | **0.944** |

**Supplementary Table 14.** Confusion matrices for the performance comparison of duplicated-region copy number callers on experimentally validated AMY1A/B/C samples. These matrices correspond to the precision and recall values reported in Supplementary Table 13

| | | | **Ground Truth** | | |
|---|---|---|---|---|---|
| **GENE** | **TOOL** | **Predicted** | NO CALL | DUP | DEL |
| AMY1A/B/C | PARASCOPY | NO CALL | 9 | 1 | 0 |
| | | DUP | 0 | 4 | 0 |
| | | DEL | 1 | 0 | 5 |
| | QUICKMER2 | NO CALL | 2 | 0 | 0 |
| | | DUP | 0 | 4 | 0 |
| | | DEL | 8 | 1 | 5 |
| | CHALLENGER$^{GENE}$ | NO CALL | 10 | 1 | 0 |
| | | DUP | 0 | 4 | 0 |
| | | DEL | 0 | 0 | 5 |

**Supplementary Table 15.** Performance comparison of duplicated-region copy number callers on MLPA-validated SMN1 samples. Bold values indicate the best performance in each metric.

| GENE | TOOLS | DEL Prec | DUP Prec | Overall Prec | DEL Rec | DUP Rec | Overall Rec | DEL F1 | DUP F1 | Overall F1 |
|---|---|---|---|---|---|---|---|---|---|---|
| SMN1 | PARASCOPY | 0.000 | 0.000 | 0.000 | 0.000 | 0.000 | 0.000 | 0.000 | 0.000 | 0.000 |
| | QUICKMER2 | 0.273 | **1.000** | 0.636 | **1.000** | 0.333 | 0.667 | 0.429 | 0.500 | 0.464 |
| | SMNCopyNumberCaller | **1.000** | **1.000** | **1.000** | **1.000** | **1.000** | **1.000** | **1.000** | **1.000** | **1.000** |
| | CHALLENGER$^{GENE}$ | **1.000** | 0.857 | 0.929 | 0.667 | **1.000** | 0.833 | 0.800 | 0.923 | 0.862 |

**Supplementary Table 16.** Confusion matrices demonstrating the performance of duplicated-region copy number callers on MLPA-validated SMN1 samples. These matrices correspond to the precision and recall metrics shown in Supplementary Table in 15

| | | | Ground Truth | | |
|---|---|---|---|---|---|
| **GENE** | **TOOL** | **Predicted** | NO CALL | DUP | DEL |
| SMN1 | PARASCOPY | NO CALL | 12 | 6 | 3 |
| | | DUP | 0 | 0 | 0 |
| | | DEL | 0 | 0 | 0 |
| | QUICKMER2 | NO CALL | 4 | 4 | 0 |
| | | DUP | 0 | 2 | 0 |
| | | DEL | 8 | 0 | 3 |
| | SMNCopyNumberCaller | NO CALL | 12 | 0 | 0 |
| | | DUP | 0 | 6 | 0 |
| | | DEL | 0 | 0 | 3 |
| | CHALLENGER$^{GENE}$ | NO CALL | 11 | 0 | 1 |
| | | DUP | 1 | 6 | 0 |
| | | DEL | 0 | 0 | 2 |

**Supplementary Table 17.** Performance comparison of duplicated-region copy number callers on experimentally validated AMY2A samples. Bold values indicate the best performance for each metric.

| **GENE** | **TOOLS** | **DEL Prec** | **DUP Prec** | **Overall Prec** | **DEL Rec** | **DUP Rec** | **Overall Rec** | **DEL F1** | **DUP F1** | **Overall F1** |
|---|---|---|---|---|---|---|---|---|---|---|
| AMY2A | PARASCOPY | **1.000** | **1.000** | **1.000** | 0.800 | 0.333 | 0.567 | 0.889 | 0.500 | 0.694 |
| | QUICKMER2 | **1.000** | **1.000** | **1.000** | **1.000** | **1.000** | **1.000** | **1.000** | **1.000** | **1.000** |
| | CHALLENGER$^{GENE}$ | **1.000** | **1.000** | **1.000** | **1.000** | **1.000** | **1.000** | **1.000** | **1.000** | **1.000** |

**Supplementary Table 18.** Confusion matrices illustrating the performance of duplicated-region copy number callers on experimentally validated AMY2A samples. These matrices correspond to the precision and recall metrics summarized in Supplementary Table 17

| | | | Ground Truth | | |
|---|---|---|---|---|---|
| **GENE** | **TOOL** | **Predicted** | NO CALL | DUP | DEL |
| AMY2A | PARASCOPY | NO CALL | 11 | 4 | 1 |
| | | DUP | 0 | 2 | 0 |
| | | DEL | 0 | 0 | 4 |
| | QUICKMER2 | NO CALL | 11 | 0 | 0 |
| | | DUP | 0 | 6 | 0 |
| | | DEL | 0 | 0 | 5 |
| | CHALLENGER$^{GENE}$ | NO CALL | 11 | 0 | 0 |
| | | DUP | 0 | 6 | 0 |
| | | DEL | 0 | 0 | 5 |

**Supplementary Table 19.** Performance comparison of duplicated-region copy number callers on MLPA-validated SMN2 samples. Bold values indicate the best performance across metrics.

| **GENE** | **TOOLS** | **DEL Prec** | **DUP Prec** | **Overall Prec** | **DEL Rec** | **DUP Rec** | **Overall Rec** | **DEL F1** | **DUP F1** | **Overall F1** |
|---|---|---|---|---|---|---|---|---|---|---|
| SMN2 | PARASCOPY | 0.000 | 0.000 | 0.000 | 0.000 | 0.000 | 0.000 | 0.000 | 0.000 | 0.000 |
| | QUICKMER2 | **1.000** | 0.429 | 0.714 | 0.333 | **1.000** | 0.667 | 0.500 | 0.600 | 0.550 |
| | SMNCopyNumberCaller | **1.000** | **1.000** | **1.000** | **1.000** | **1.000** | **1.000** | **1.000** | **1.000** | **1.000** |
| | CHALLENGER$^{GENE}$ | **1.000** | **1.000** | **1.000** | **1.000** | **1.000** | **1.000** | **1.000** | **1.000** | **1.000** |

**Supplementary Table 20.** Confusion matrices representing the performance of duplicated-region copy number callers on MLPA-validated SMN2 samples. These matrices correspond to the precision and recall values summarized in Supplementary Table in 19

| | | | Ground Truth | | |
|---|---|---|---|---|---|
| **GENE** | **TOOL** | **Predicted** | NO CALL | DUP | DEL |
| SMN2 | PARASCOPY | NO CALL | 12 | 6 | 6 |
| | | DUP | 0 | 0 | 0 |
| | | DEL | 0 | 0 | 0 |
| | QUICKMER2 | NO CALL | 4 | 0 | 4 |
| | | DUP | 8 | 6 | 0 |
| | | DEL | 0 | 0 | 2 |
| | SMNCopyNumberCaller | NO CALL | 12 | 0 | 0 |
| | | DUP | 0 | 6 | 0 |
| | | DEL | 0 | 0 | 6 |
| | CHALLENGER$^{GENE}$ | NO CALL | 12 | 0 | 0 |
| | | DUP | 0 | 6 | 0 |
| | | DEL | 0 | 0 | 6 |

**Supplementary Table 21.** Performance comparison of duplicated-region copy number callers on MLPA-validated SMN1/2 samples. Bold values denote the best performance for each metric.

| **GENE** | **TOOLS** | **DEL Prec** | **DUP Prec** | **Overall Prec** | **DEL Rec** | **DUP Rec** | **Overall Rec** | **DEL F1** | **DUP F1** | **Overall F1** |
|---|---|---|---|---|---|---|---|---|---|---|
| SMN1/2 | PARASCOPY | **1.000** | **1.000** | **1.000** | 0.889 | **1.000** | 0.944 | 0.941 | **1.000** | 0.971 |
| | QUICKMER2 | **1.000** | **1.000** | **1.000** | **1.000** | **1.000** | **1.000** | **1.000** | **1.000** | **1.000** |
| | SMNCopyNumberCaller | **1.000** | **1.000** | **1.000** | **1.000** | **1.000** | **1.000** | **1.000** | **1.000** | **1.000** |
| | CHALLENGER$^{GENE}$ | **1.000** | **1.000** | **1.000** | **1.000** | **1.000** | **1.000** | **1.000** | **1.000** | **1.000** |

**Supplementary Table 22.** Confusion matrices displaying the performance of duplicated-region copy number callers on MLPA-validated SMN1/2 samples. These matrices correspond to the precision and recall results reported in Supplementary Table in 21

| | | | Ground Truth | | |
|---|---|---|---|---|---|
| **GENE** | **TOOL** | **Predicted** | NO CALL | DUP | DEL |
| SMN1/2 | PARASCOPY | NO CALL | 24 | 0 | 1 |
| | | DUP | 0 | 12 | 0 |
| | | DEL | 0 | 0 | 8 |
| | QUICKMER2 | NO CALL | 24 | 0 | 0 |
| | | DUP | 0 | 12 | 0 |
| | | DEL | 0 | 0 | 9 |
| | SMNCopyNumberCaller | NO CALL | 24 | 0 | 0 |
| | | DUP | 0 | 12 | 0 |
| | | DEL | 0 | 0 | 9 |
| | CHALLENGER$^{GENE}$ | NO CALL | 24 | 0 | 0 |
| | | DUP | 0 | 12 | 0 |
| | | DEL | 0 | 0 | 9 |

**Supplementary Table 23.** Ablation study of the CHALLENGER model showing the impact of removing key components on deletion, duplication, and overall F1 scores. Each variant demonstrates the contribution of the Transformer encoder, gene embeddings, and positional embeddings to the model's overall performance.

| | **DEL F1 Score** | **DUP F1 Score** | **Overall F1 Score** |
|---|---|---|---|
| CHALLENGER$^{LR}$ | **0.78** | **0.63** | **0.71** |
| CHALLENGER$^{LR}$ (w/o Transformer) | 0.64 | 0.53 | 0.58 |
| CHALLENGER$^{LR}$ (w/o Gene Embed.) | 0.40 | 0.50 | 0.45 |
| CHALLENGER$^{LR}$ (w/o Pos. Embed.) | 0.68 | 0.54 | 0.61 |

**Supplementary Table 24.** List of 550 samples from the 1000 Genomes dataset selected for pre-training CHALLENGER.

| | | | | | | | | | |
|---|---|---|---|---|---|---|---|---|---|
| HG00472 | HG01119 | HG02190 | HG01444 | HG01113 | HG00288 | HG01860 | HG00237 | HG02023 | HG02356 |
| HG02292 | HG01809 | HG00622 | HG02239 | HG01918 | HG01066 | HG01372 | HG01048 | HG01173 | HG01353 |
| HG01468 | HG00336 | HG02089 | HG01359 | HG01606 | HG01869 | HG00879 | HG00867 | HG01111 | HG01474 |
| HG02079 | HG01075 | HG01857 | HG02139 | HG02230 | HG01531 | HG00473 | HG02332 | HG01051 | HG01164 |
| HG01605 | HG01072 | HG01198 | HG01865 | HG01253 | HG01845 | HG00736 | HG01354 | HG01802 | HG01610 |
| HG01187 | HG01801 | HG01049 | HG01110 | HG01485 | HG00253 | HG01598 | HG01926 | HG01088 | HG01921 |
| HG01398 | HG02182 | HG01360 | HG02231 | HG00421 | HG00554 | HG01497 | HG01190 | HG00242 | HG00631 |
| HG01847 | HG01351 | HG01783 | HG01271 | HG02262 | HG01844 | HG00331 | HG00304 | HG02307 | HG00442 |
| HG01914 | HG01461 | HG00096 | HG01256 | HG01507 | HG01811 | HG01489 | HG02143 | HG01509 | HG00662 |
| HG01069 | HG01756 | HG01492 | HG02142 | HG00252 | HG00864 | HG01956 | HG00277 | HG00419 | HG01396 |
| HG00332 | HG01437 | HG00367 | HG01272 | HG02238 | HG00260 | HG02156 | HG00251 | HG00614 | HG02233 |
| HG02075 | HG01880 | HG01958 | HG01086 | HG01063 | HG01628 | HG01840 | HG01241 | HG00590 | HG00674 |
| HG02069 | HG00844 | HG00268 | HG01047 | HG01571 | HG01341 | HG01205 | HG00326 | HG00632 | HG01055 |
| HG00250 | HG02113 | HG01870 | HG01768 | HG02072 | HG02186 | HG01085 | HG01684 | HG00099 | HG01124 |
| HG00583 | HG01858 | HG01572 | HG01438 | HG00422 | HG01747 | HG02090 | HG02298 | HG00325 | HG02166 |
| HG01623 | HG00478 | HG00436 | HG00543 | HG01257 | HG01149 | HG01920 | HG01174 | HG01441 | HG00640 |
| HG00278 | HG02317 | HG01710 | HG02259 | HG02061 | HG00449 | HG02084 | HG02028 | HG02006 | HG01046 |
| HG01200 | HG01139 | HG02009 | HG01350 | HG02085 | HG01894 | HG00464 | HG01583 | HG02284 | HG02019 |
| HG02291 | HG02215 | HG01125 | HG01402 | HG01848 | HG01095 | HG02282 | HG01518 | HG01079 | HG02017 |
| HG00334 | HG00309 | HG01953 | HG01133 | HG01284 | HG01597 | HG01501 | HG00310 | HG02322 | HG02047 |
| HG01630 | HG01204 | HG00382 | HG00372 | HG00653 | HG01915 | HG01326 | HG00353 | HG01512 | HG01369 |
| HG02285 | HG00245 | HG00551 | HG00592 | HG00634 | HG00276 | HG01248 | HG00692 | HG00280 | HG02337 |
| HG01893 | HG00537 | HG02081 | HG01494 | HG01777 | HG02140 | HG02051 | HG01889 | HG01440 | HG00607 |
| HG01790 | HG01140 | HG01805 | HG01521 | HG00536 | HG01699 | HG01176 | HG01323 | HG01362 | HG01816 |
| HG01851 | HG01625 | HG01631 | HG00542 | HG02070 | HG00598 | HG00595 | HG00463 | HG02050 | HG00239 |
| HG01412 | HG02188 | HG02064 | HG02278 | HG00256 | HG02088 | HG01191 | HG00324 | HG01992 | HG01971 |
| HG00663 | HG00375 | HG00327 | HG01762 | HG00534 | HG01708 | HG01890 | HG01771 | HG00560 | HG02016 |
| HG02082 | HG01305 | HG01668 | HG00445 | HG00384 | HG02179 | HG02185 | HG01134 | HG01935 | HG02057 |
| HG00448 | HG01613 | HG00361 | HG01912 | HG00345 | HG00369 | HG01148 | HG01524 | HG01593 | HG01990 |
| HG02151 | HG01991 | HG01058 | HG01551 | HG01694 | HG01137 | HG00357 | HG01413 | HG02147 | HG01879 |
| HG01883 | HG00531 | HG01131 | HG01927 | HG01988 | HG00257 | HG02223 | HG02283 | HG01866 | HG02353 |
| HG01941 | HG01686 | HG02232 | HG00254 | HG01171 | HG01578 | HG00623 | HG00376 | HG01259 | HG01797 |
| HG00321 | HG01974 | HG00732 | HG00672 | HG02138 | HG01702 | HG02054 | HG00263 | HG00360 | HG01950 |
| HG02277 | HG01980 | HG00329 | HG01308 | HG02343 | HG01516 | HG02309 | HG01101 | HG01620 | HG00479 |
| HG01073 | HG01393 | HG00404 | HG00306 | HG02219 | HG02031 | HG01603 | HG02136 | HG02104 | HG01938 |
| HG01789 | HG01488 | HG02058 | HG00328 | HG01456 | HG01281 | HG00701 | HG00619 | HG00475 | HG00625 |
| HG00271 | HG00378 | HG01348 | HG00368 | HG01673 | HG00629 | HG01432 | HG01932 | HG00656 | HG02102 |
| HG01600 | HG01595 | HG01112 | HG00362 | HG00476 | HG00265 | HG01612 | HG00699 | HG01682 | HG01849 |
| HG02299 | HG01976 | HG01817 | HG00246 | HG00236 | HG01334 | HG02165 | HG00525 | HG01356 | HG01530 |
| HG01061 | HG01800 | HG01810 | HG01108 | HG01345 | HG00338 | HG01363 | HG00315 | HG00234 | HG00690 |
| HG02272 | HG01779 | HG00410 | HG02351 | HG00596 | HG00318 | HG00452 | HG01504 | HG02060 | HG01757 |
| HG00743 | HG01269 | HG01250 | HG01565 | HG01589 | HG01121 | HG00500 | HG01704 | HG00599 | HG02134 |
| HG00232 | HG02312 | HG01525 | HG02086 | HG01162 | HG01806 | HG01871 | HG02116 | HG01550 | HG01566 |
| HG01855 | HG00740 | HG01965 | HG00684 | HG01945 | HG00978 | HG02152 | HG01617 | HG01031 | HG01197 |
| HG01670 | HG01130 | HG01375 | HG01979 | HG01462 | HG01342 | HG02153 | HG00610 | HG01776 | HG00341 |
| HG01970 | HG01455 | HG00255 | HG02266 | HG00379 | HG01495 | HG01815 | HG01384 | HG02256 | HG00355 |
| HG01459 | HG00628 | HG00956 | HG01029 | HG01182 | HG02220 | HG01798 | HG01168 | HG02330 | HG00356 |
| HG02286 | HG00346 | HG01812 | HG02315 | HG01503 | HG01092 | HG00313 | HG02155 | HG01924 | HG00657 |
| HG01968 | HG01344 | HG00350 | HG01794 | HG01161 | HG00766 | HG00611 | HG02073 | HG01951 | HG02164 |
| HG01142 | HG00409 | HG00403 | HG00698 | HG02053 | HG00238 | HG00581 | HG01107 | HG02187 | HG01765 |
| HG01859 | HG01882 | HG01868 | HG00351 | HG01506 | HG02128 | HG01537 | HG02040 | HG01357 | HG00281 |
| HG02250 | HG01302 | HG00428 | HG00593 | HG00290 | HG02137 | HG02325 | HG01105 | HG01077 | HG00734 |
| HG01491 | HG02314 | HG02002 | HG00566 | HG01247 | HG01709 | HG02095 | HG02048 | HG00689 | HG02265 |
| HG00330 | HG00358 | HG02076 | HG01447 | HG00097 | HG01556 | HG00261 | HG01136 | HG00373 | HG01365 |
| HG01094 | HG01863 | HG01104 | HG01700 | HG01936 | HG02345 | HG01366 | HG00443 | HG00337 | HG01389 |

**Supplementary Table 25.** List of 230 samples from the 1000 Genomes dataset used to fine-tune the model CHALLENGER$^{LR}$.

| | | | | | | | | | |
|---|---|---|---|---|---|---|---|---|---|
| HG00357 | HG01617 | HG02284 | HG00436 | HG01341 | HG00360 | HG02182 | HG00844 | HG02223 | HG01673 |
| HG01694 | HG02332 | HG02309 | HG02250 | HG01131 | HG02095 | HG00610 | HG02009 | HG02089 | HG00278 |
| HG02285 | HG00321 | HG02136 | HG00261 | HG00368 | HG01396 | HG00313 | HG01953 | HG00325 | HG01598 |
| HG01168 | HG01133 | HG01756 | HG01389 | HG02239 | HG01072 | HG01148 | HG01918 | HG01779 | HG02128 |
| HG01765 | HG01197 | HG02058 | HG01578 | HG01256 | HG02139 | HG01187 | HG01951 | HG02337 | HG01606 |
| HG01595 | HG01058 | HG02006 | HG01134 | HG01362 | HG01589 | HG00346 | HG01776 | HG00472 | HG02152 |
| HG00537 | HG01073 | HG01356 | HG01474 | HG01700 | HG01600 | HG02079 | HG02238 | HG01684 | HG01805 |
| HG01302 | HG01121 | HG00452 | HG02153 | HG01935 | HG00324 | HG02231 | HG00271 | HG00338 | HG01851 |
| HG01107 | HG00634 | HG02265 | HG01583 | HG01048 | HG00099 | HG01597 | HG01800 | HG02277 | HG00378 |
| HG01047 | HG00475 | HG01699 | HG02072 | HG02291 | HG01879 | HG01806 | HG01136 | HG00619 | HG01894 |
| HG01870 | HG00443 | HG01509 | HG01492 | HG01932 | HG01384 | HG02307 | HG01108 | HG00254 | HG01566 |
| HG01571 | HG02031 | HG01344 | HG00096 | HG01798 | HG00583 | HG00268 | HG02165 | HG01174 | HG01789 |
| HG01494 | HG01883 | HG01412 | HG00657 | HG01950 | HG01525 | HG02143 | HG01259 | HG01551 | HG01130 |
| HG01817 | HG01366 | HG00419 | HG00252 | HG01630 | HG01550 | HG01066 | HG01086 | HG01988 | HG01504 |
| HG02164 | HG01055 | HG00632 | HG01375 | HG02325 | HG00442 | HG01518 | HG00361 | HG01489 | HG01506 |
| HG01241 | HG02064 | HG02016 | HG01124 | HG01269 | HG02082 | HG02069 | HG01865 | HG01512 | HG01605 |
| HG01085 | HG01142 | HG01119 | HG01866 | HG00551 | HG01398 | HG00479 | HG01507 | HG01455 | HG02292 |
| HG00622 | HG01612 | HG00255 | HG01345 | HG00740 | HG01871 | HG02155 | HG01992 | HG00336 | HG01572 |
| HG01284 | HG01447 | HG01971 | HG01915 | HG01809 | HG02017 | HG00684 | HG00464 | HG02283 | HG02060 |
| HG01815 | HG00315 | HG00367 | HG00692 | HG01840 | HG01308 | HG01965 | HG01198 | HG01945 | HG01257 |
| HG00596 | HG00242 | HG00766 | HG01530 | HG01990 | HG00409 | HG01101 | HG02266 | HG00449 | HG00334 |
| HG01991 | HG02137 | HG00736 | HG02138 | HG00607 | HG02081 | HG01348 | HG01890 | HG00566 | HG02156 |
| HG01794 | HG02053 | HG01880 | HG01863 | HG00404 | HG01859 | HG02314 | HG02116 | HG01104 | HG01137 |

**Supplementary Table 26.** List of 30 samples from the 1000 Genomes dataset used to evaluate the performance of CHALLENGER$^{LR}$.

| | | | | | | | | | |
|---|---|---|---|---|---|---|---|---|---|
| HG02013 | HG01060 | HG00269 | HG01917 | HG01766 | HG00513 | HG00335 | HG00311 | HG02144 | HG02318 |
| HG00650 | HG01167 | HG02146 | HG01311 | HG02067 | HG02130 | HG01596 | HG02133 | HG01948 | HG00651 |
| HG00739 | HG01961 | HG02274 | HG00584 | HG02180 | HG01054 | HG00626 | HG01377 | HG02107 | HG02087 |

**Supplementary Table 27.** List of samples from the 1000 Genomes dataset used to fine-tune and test the performance of CHALLENGER$^{EXPERT}$.

| **Fine-Tuning Samples** | **Test Samples** |
|---|---|
| HG00513 | HG00731 |
| HG00732 | NA19238 |
| HG00733 | NA19240 |
| NA19239 | - |
| HG00512 | - |
| HG00514 | - |

**Supplementary Table 28.** List of samples from the 1000 Genomes dataset used to fine-tune and test the performance on SMN genes.

| | Fine-Tuning Samples | | Test Samples | |
|---|---|---|---|---|
| **Gene Name** | **Sample Name** | **CNV** | **Sample Name** | **CNV** |
| SMN1 | HG01948 | DEL | HG01612 | DEL |
| | HG00324 | DEL | HG03928 | DEL |
| | NA20812 | DEL | NA20764 | DEL |
| | HG01684 | DUP | HG03603 | DUP |
| | HG02231 | DUP | HG00179 | DUP |
| | HG02089 | DUP | HG00148 | DUP |
| | HG00566 | DUP | HG03473 | DUP |
| | NA20798 | DUP | HG01500 | DUP |
| | HG00327 | NO-CALL | HG03124 | DUP |
| | HG00421 | NO-CALL | HG00464 | NO-CALL |
| | HG00261 | NO-CALL | HG01997 | NO-CALL |
| | HG02239 | NO-CALL | HG01685 | NO-CALL |
| | HG02232 | NO-CALL | HG00524 | NO-CALL |
| | HG01992 | NO-CALL | HG00451 | NO-CALL |
| | HG01935 | NO-CALL | HG02221 | NO-CALL |
| | HG01694 | NO-CALL | HG02008 | NO-CALL |
| | HG01979 | NO-CALL | HG01923 | NO-CALL |
| | HG01980 | NO-CALL | HG00559 | NO-CALL |
| | HG02147 | NO-CALL | HG00371 | NO-CALL |
| | HG01678 | NO-CALL | HG02146 | NO-CALL |
| | - | - | HG00285 | NO-CALL |
| SMN2 | HG00326 | DEL | HG02002 | DEL |
| | HG00475 | DEL | HG01954 | DEL |
| | HG00319 | DEL | HG01953 | DEL |
| | HG02090 | DEL | HG02238 | DEL |
| | HG02259 | DEL | HG02220 | DEL |
| | HG01933 | DEL | HG00449 | DEL |
| | HG01747 | DUP | HG00233 | DUP |
| | HG01971 | DUP | HG00109 | DUP |
| | HG00626 | DUP | HG01515 | DUP |
| | HG00318 | DUP | HG00693 | DUP |
| | HG00329 | DUP | HG02814 | DUP |
| | HG01260 | DUP | HG03616 | DUP |
| | HG00353 | NO-CALL | HG02233 | NO-CALL |
| | HG01776 | NO-CALL | HG01374 | NO-CALL |
| | HG01777 | NO-CALL | HG00580 | NO-CALL |
| | HG01756 | NO-CALL | HG00284 | NO-CALL |
| | HG01699 | NO-CALL | HG00557 | NO-CALL |
| | HG01771 | NO-CALL | HG00651 | NO-CALL |
| | HG01682 | NO-CALL | HG01602 | NO-CALL |
| | HG01945 | NO-CALL | HG01254 | NO-CALL |
| | HG01921 | NO-CALL | HG00650 | NO-CALL |
| | HG01950 | NO-CALL | HG01277 | NO-CALL |
| | HG01976 | NO-CALL | HG00708 | NO-CALL |
| | HG00513 | NO-CALL | HG01618 | NO-CALL |

**Supplementary Table 29.** List of samples from the 1000 Genomes dataset used to fine-tune and test the performance on AMY1A/B/C and AMY2A genes.

| | **Fine-Tuning Samples** | | **Test Samples** | |
|---|---|---|---|---|
| **Gene Name** | **Sample Name** | **CNV** | **Sample Name** | **CNV** |
| AMY2A | NA19005 | NO-CALL | NA18995 | NO-CALL |
| | NA18976 | NO-CALL | NA18998 | NO-CALL |
| | NA19201 | NO-CALL | NA18999 | NO-CALL |
| | NA12761 | NO-CALL | NA18953 | NO-CALL |
| | NA18633 | NO-CALL | NA18956 | NO-CALL |
| | NA18522 | NO-CALL | NA18952 | NO-CALL |
| | NA18948 | NO-CALL | NA18980 | NO-CALL |
| | NA18949 | NO-CALL | NA18951 | NO-CALL |
| | NA18940 | NO-CALL | NA18968 | NO-CALL |
| | NA18959 | NO-CALL | NA18981 | NO-CALL |
| | NA12275 | NO-CALL | NA18960 | NO-CALL |
| | NA18961 | NO-CALL | NA18624 | DEL |
| | NA12156 | DEL | NA07034 | DEL |
| | NA12815 | DEL | NA06985 | DEL |
| | NA11829 | DEL | NA07045 | DEL |
| | NA11995 | DEL | NA12343 | DEL |
| | NA11992 | DEL | NA12873 | DUP |
| | NA12146 | DEL | NA12043 | DUP |
| | NA19204 | DUP | NA12239 | DUP |
| | NA19012 | DUP | NA19222 | DUP |
| | NA12155 | DUP | NA19223 | DUP |
| | NA12763 | DUP | NA19200 | DUP |
| | NA11830 | DUP | - | - |
| | NA11881 | DUP | - | - |
| AMY1A/B/C | NA19201 | NO-CALL | NA12751 | NO-CALL |
| | NA18959 | NO-CALL | NA12762 | NO-CALL |
| | NA18961 | NO-CALL | NA12812 | NO-CALL |
| | NA18999 | NO-CALL | NA12814 | NO-CALL |
| | NA18951 | NO-CALL | NA07056 | NO-CALL |
| | NA18981 | NO-CALL | NA11839 | NO-CALL |
| | NA18967 | NO-CALL | NA12003 | NO-CALL |
| | NA18973 | NO-CALL | NA12044 | NO-CALL |
| | NA18970 | NO-CALL | NA12144 | NO-CALL |
| | NA18966 | NO-CALL | NA12154 | NO-CALL |
| | NA11840 | NO-CALL | NA18522 | DEL |
| | NA12750 | NO-CALL | NA18945 | DEL |
| | NA18633 | DEL | NA12760 | DEL |
| | NA18969 | DEL | NA11831 | DEL |
| | NA12156 | DEL | NA11994 | DEL |
| | NA11829 | DEL | NA18976 | DUP |
| | NA11832 | DEL | NA18949 | DUP |
| | NA12239 | DEL | NA12275 | DUP |
| | NA12813 | DEL | NA18956 | DUP |
| | NA19204 | DUP | NA18952 | DUP |
| | NA18974 | DUP | - | - |
| | NA18992 | DUP | - | - |
| | NA19239 | DUP | - | - |
| | NA12763 | DUP | - | - |
| | NA11995 | DUP | - | - |
| | NA18953 | DUP | - | - |

**Supplementary Table 30.** List of samples from the 1000 Genomes dataset used to fine-tune and test the performance on NPY4RA/B genes.

| | **Fine-Tuning Samples** | | **Test Samples** | |
|---|---|---|---|---|
| **Gene Name** | **Sample Name** | **CNV** | **Sample Name** | **CNV** |
| NPY4RA/B | NA18961 | NO-CALL | NA10847 | NO-CALL |
| | NA12717 | NO-CALL | NA18504 | NO-CALL |
| | NA18542 | NO-CALL | NA18627 | NO-CALL |
| | NA18603 | NO-CALL | NA18745 | DUP |
| | NA18948 | DUP | NA18536 | DUP |
| | NA18949 | DUP | NA18510 | DUP |
| | NA18959 | DUP | NA18519 | DUP |
| | NA12155 | DUP | NA10851 | DUP |
| | NA18517 | DUP | NA18940 | DUP |

**Supplementary Table 31.** List of genes without deletion events in the training/fine-tuning set but with deletion events in the test set, for which CHALLENGER achieved a 100% F1 score.

| | | | | |
|---|---|---|---|---|
| AIFM3 | AK5 | ARFGEF3 | C2orf80 | C3orf84 |
| CRKL | ENSG00000288000 | FER1L6 | FIG4 | FRZB |
| IWS1 | LRRC40 | LRRC74B | LZTR1 | MCFD2 |
| NCALD | P2RX6 | PARD3 | PDGFC | PHF3 |
| REC114 | RNF4 | SERPIND1 | SH3GL2 | SLC13A4 |
| SLC7A4 | SNAP29 | SNX4 | THAP7 | TMEM266 |
| TTC7A | VTI1A | WHRN | ZBTB12 | |

**Supplementary Table 32.** List of genes without duplication events in the training/fine-tuning set but with duplication events in the test set, for which CHALLENGER achieved a 100% F1 score.

| | | | | |
|---|---|---|---|---|
| ABHD12 | ADNP2 | AQR | BTNL8 | ENSG00000288711 |
| FOXL2 | NDUFAF2 | PACSIN2 | PARD6G | PILRA |
| PILRB | SAMD5 | STON1 | STON1-GTF2A1L | SULF1 |
| TMEM14C | | | | |

**Supplementary Table 33.** List of genes with deletion allele frequency below %1 in the training/fine-tuning set, for which CHALLENGER achieves a 100% F1 score.

| | | | | |
|---|---|---|---|---|
| ACER3 | ACOT2 | AIFM3 | AK5 | ALG1 |
| ARFGEF3 | ARHGAP8 | BICD1 | C16orf89 | C2orf80 |
| C3orf84 | C6 | CADPS | CDC42BPA | CFAP20DC |
| CFAP61 | CHN1 | CHRNB3 | CLN8 | CNIH3 |
| CPA6 | CPVL | CRADD | CRKL | CYP1B1 |
| DHRS7C | DOCK3 | EGF | ENSG00000196826 | ENSG00000284461 |
| ENSG00000284931 | ENSG00000285082 | ENSG00000288000 | ERC1 | FAM174B |
| FER1L6 | FGF1 | FIG4 | FMNL2 | FRZB |
| GABRB1 | GARNL3 | GLRB | GLT1D1 | GLYAT |
| GNAL | GPR137B | GPR137C | GPR161 | GRHL2 |
| GRM3 | GTF2F2 | HBS1L | HECW1 | INSR |
| IWS1 | KDM2A | KIAA1217 | KIF1B | KRTAP4-11 |
| KRTAP4-9 | LMNTD1 | LRRC40 | LRRC74B | LZTR1 |
| MAP2K5 | MAP3K13 | MCFD2 | MRC1 | MTHFS |
| NACC2 | NCALD | NCMAP | NECTIN3 | NEDD4L |
| NOL10 | OR3A2 | OR4F15 | OR4F6 | OR9Q1 |
| OXR1 | P2RX6 | PAK5 | PALM2AKAP2 | PANX1 |
| PARD3 | PCNT | PDGFC | PDSS2 | PHF3 |
| PI4KA | PIK3AP1 | PLA2G2C | PLCL2 | PLEKHA7 |
| PPP6R3 | PRKCE | PTBP3 | PTPRO | RAB11FIP3 |
| RAB3GAP2 | RC3H1 | REC114 | RERGL | RNF4 |
| ROCK2 | RPS6KC1 | RUNX1 | SCAF8 | SCN9A |
| SERPIND1 | SH3GL2 | SLC13A4 | SLC1A1 | SLC22A2 |
| SLC7A4 | SNAP29 | SNX4 | SRGAP3 | SSH2 |
| ST20-MTHFS | STAT4 | SYT16 | TBX15 | TCAIM |
| TCF12 | TFEC | THAP7 | TMEFF2 | TMEM132C |
| TMEM266 | TSC22D1 | TTC7A | UGT1A3 | UGT1A4 |
| UGT1A5 | UGT1A6 | UGT1A7 | UGT1A9 | USP32 |
| VRK2 | VTI1A | VWA3B | WDR76 | WHRN |
| WNT2B | ZBTB12 | ZNF264 | ZNF709 | |

**Supplementary Table 34.** List of genes with duplication allele frequency below %1 in the training/fine-tuning set, for which CHALLENGER achieves a 100% F1 score.

| | | | | |
|---|---|---|---|---|
| ABHD12 | ADNP2 | AQR | ATAD3B | ATAD3C |
| BTNL3 | BTNL8 | CDH1 | CDH18 | CFAP20DC |
| DNASE1 | DOP1B | ENSG00000268870 | ENSG00000288711 | FOXL2 |
| HAL | HVCN1 | LTA4H | MYO7B | NANOGNB |
| NDUFAF2 | PACSIN2 | PARD6G | PCDHGA1 | PCDHGA10 |
| PCDHGB6 | PCDHGB7 | PILRA | PILRB | RASSF3 |
| RBM28 | RUNX1 | SAMD5 | STON1 | STON1-GTF2A1L |
| SULF1 | TMEM14C | TRIM77 | TTC28 | VKORC1L1 |

#### Supplementary References
